# IMU-based upper limb kinematics validation: sensor fusion optimisation, sensor-to-segment alignment, and joint angle accuracy

**DOI:** 10.64898/2026.09.29.755510

**Authors:** Arthur Fabre, Graham Kerr, Peter Pivonka, Wolbert van den Hoorn

## Abstract

Conventional clinical assessment of upper limb kinematics relies on goniometry or visual estimation, methods that provide limited information on multi-segment 3D kinematics and can have poor inter-rater reliability. Inertial measurement units (IMUs) represent a viable alternative; however, extracting anatomically meaningful joint angles from skin-mounted sensors remains challenging owing to sensor-to-segment misalignment, soft-tissue artefact, and suboptimal orientation estimation under dynamic conditions. This study developed and validated a multi-segment IMU-based framework for upper limb kinematic assessment, addressing three stages of the measurement pipeline: (i) sensor fusion gain optimisation as a function of task mean segment angular velocity; (ii) accuracy of an anatomical landmark registration-based alignment method; and (iii) comparative evaluation of landmark-based alignment against manual and neutral-pose alignment approaches. Twenty participants with typical shoulder function performed clinical and functional upper limb tasks, recorded simultaneously with an optoelectronic motion capture system and six IMUs tracking the thorax, clavicle, scapula, humerus, forearm, and hand. Orientation estimation used a gradient-descent complementary filter with velocity-dependent gain modulation. Sensor-to-segment alignment was achieved via manual, neutral-pose, and landmark registration methods. Mean sensor orientation errors ranged from 1.5° to 3° across segments/tasks and increased with angular velocity. Landmark registration produced substantially lower joint angle errors than manual and neutral-pose alignment across all joints (root mean square error range: 3.7°–7.8°; manual: 8.3°–24.2°; neutral: 15.5°−35.5°). These results indicate that IMU-derived kinematic accuracy is context-dependent, varying with joint and movement type, suggesting that accuracy should be established for the specific movements and joints relevant to each clinical application.

## 1. Introduction

Accurate assessment of upper limb function is important for tracking outcomes after surgery or rehabilitation. Routine clinical assessments, however, rely on goniometers or visual estimations, often lacking sensitivity (Donnelly et al., 2013) and reliability (Hayes et al., 2001) and reduce shoulder function to gross thoracohumeral angles. Wearable inertial measurement units (IMUs) offer an objective alternative for 3D kinematic assessment (Stefana et al., 2021). Achieving this requires IMU-derived joint angles to be anatomically meaningful and sufficiently accurate across relevant joints and tasks.

Extracting anatomically meaningful joint angles from skin-mounted IMUs is challenging because raw sensor orientation must be estimated in the presence of movement-related accelerations and magnetic field disturbances, and expressed in anatomical coordinate systems for each underlying body segment. Sensor-fusion algorithms must balance gyroscope integration drift with corrections for gravity and Earth magnetic fields, and accuracy changes with movement dynamics, requiring careful tuning of fusion parameters. In contrast to the commonly used Madgwick filter (Madgwick et al., 2011) which uses a single correction gain, Riddick et al. (2023) estimated IMU orientation using a gradient descent-based filter with separate gyroscope, accelerometer, and magnetometer gains within a prediction-correction framework.

Sensor-to-segment alignment is a primary source of error, particularly when bony landmarks are difficult to identify or soft tissue artefact (STA) is substantial (Zhu et al., 2023). Common approaches include manual IMU alignment, neutral-pose alignment, and functional movement-based calibrations (Bouvier et al., 2015; Hindle et al., 2021), but these can be sensitive to body morphology and participants’ ability to adopt prescribed postures or movements. Alternatively, anatomical landmarks registration using external tools may provide a more standardised link to International Society of Biomechanics (ISB) coordinate systems (Wu et al., 2005). However, its accuracy across the shoulder girdle and wrist during functional and multiplane tasks remains limited (Picerno et al., 2019).

This study aimed to develop and determine the accuracy of an IMU-based method for estimating anatomically meaningful upper-limb joint angles, using optical motion capture (OMC) as reference across three pipeline stages: IMU orientation estimation, sensor-to-segment alignment, and joint angle estimation. Specifically, we quantified IMU orientation error after speed-dependent tuning of sensor fusion gains; compared landmark-based registration with manual and neutral-pose alignment; and assessed how these alignment approaches affected joint angle errors across joints and tasks. We expected landmark registration to produce joint angle estimates closer to OMC than manual alignment and greater anatomical interpretability than neutral-pose alignment.

## 2. Methods

### 2.1. Participants

Twenty participants (10 females, 10 males; mean [SD]: age 29.7 [6.2] years, height 1.69 [0.09] m, body mass 67.0 [13.8] kg; limb dominance: 17 right, 3 left) were recruited from the local community by convenience sampling. Eligibility criteria included age ≥18 years and no upper-limb pain, history of musculoskeletal injury, or surgery. Ethical approval was granted by the Institutional Human Ethics Committee (ID 8094), and all participants provided written informed consent prior to participation.

### 2.2. Experimental setup

Upper limb movements were recorded using a twelve-camera OMC system and six IMUs (Vantage V5 and Blue Trident, VICON®). OMC and IMU data were sampled at 125 and 1125 samples/s, respectively, using Vicon Nexus and hardware-synchronised via Vicon Beacon. The Vicon reference frame was defined with the X-, Y-, and Z-axes oriented along the anterior, leftward, and superior directions of the participant, respectively. As IMU orientation estimates were expressed in the Earth/Global reference frame, the Vicon frame was aligned offline to the IMU Global frame, with the X-axis aligned to magnetic north (Appendix A.3 and A.5).

#### 2.2.1. Markers and cluster setup

Reflective markers (14 mm diameter) were attached to the skin over anatomical landmarks using double-sided tape, following ISB recommendations (Wu et al., 2005) (Figure 1). For the eight participants in whom the xiphoid process was inaccessible, a custom pointer registered this landmark relative to the thoracic marker cluster. The glenohumeral joint centre was estimated separately for the OMC and IMU system from the same humerothoracic circumduction task (performed below 90° thoracohumeral elevation). Its location was determined using the symmetrical centre of rotation estimation method (Ehrig et al., 2006) for OMC, and the symmetrical sphere-fit centre method (Crabolu et al., 2017) for IMUs, with results registered to both the scapula and humerus marker clusters and IMUs, respectively (Appendix B.2).

**Figure 1.**
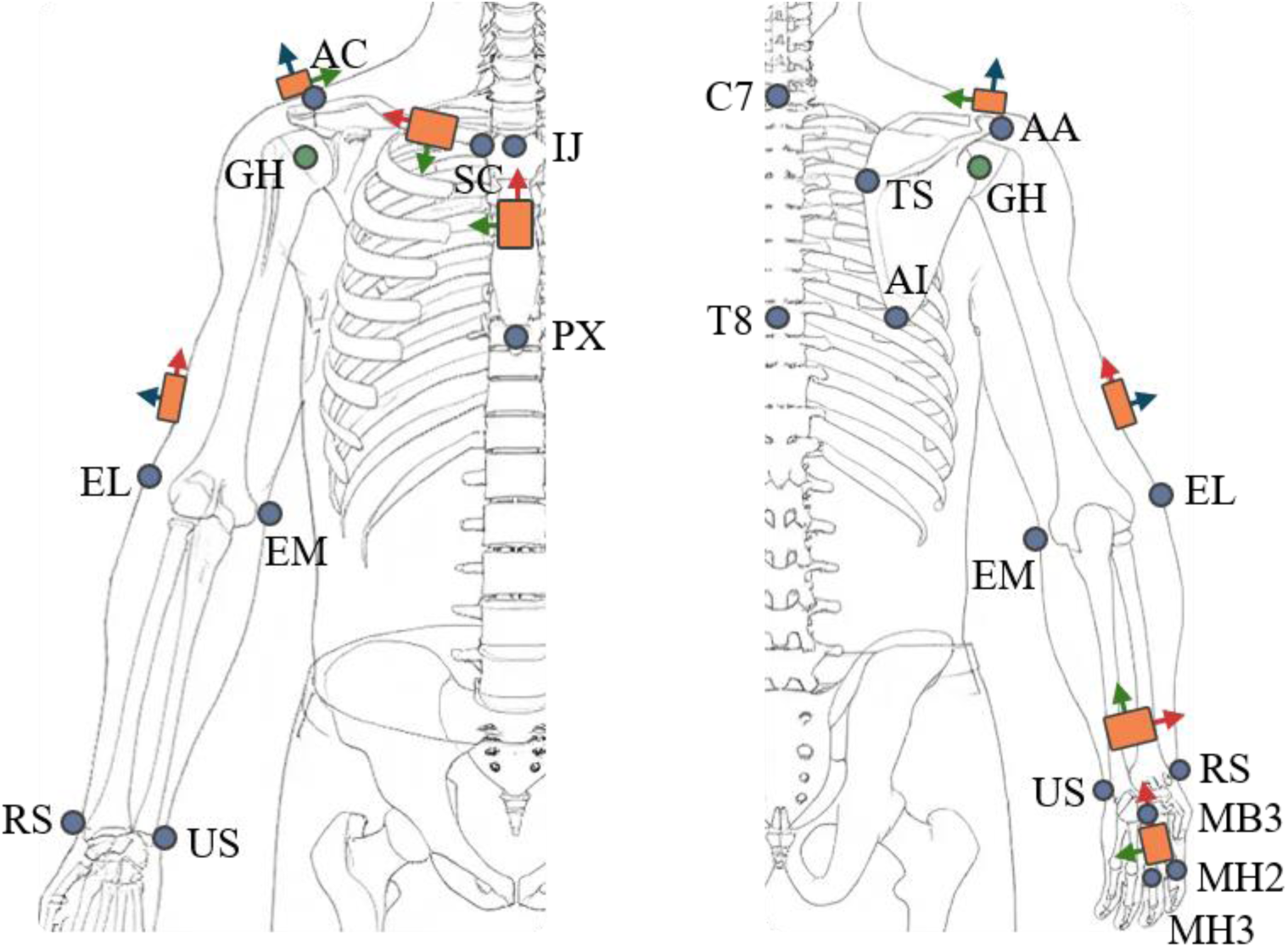
Anatomical placement of IMUs and OMC markers. Legend: Left: anterior view, Right: posterior view, Blue circles: OMC markers; Green circles: virtual glenohumeral joint rotation centre; Orange rectangles: IMUs and clusters, PX: processus xiphoideus, IJ: incisura jugularis, C7: 7th cervical vertebra, T8: 8th thoracic vertebra, SC: sternoclavicular joint, AC: acromioclavicular joint, AA: angulus acromialis, TS: trigonum spinae, AI: angulus inferior, GH: glenohumeral join rotation centre, EL: lateral epicondyle, EM: medial epicondyle, RS: radius styloid, US: ulnar styloid, MB2: 2nd metacarpal base, MH2: 2nd metacarpal head, MH3: 3rd metacarpal head, red, green and blue axes represent the local x, y and z axes of IMUs, respectively.

#### 2.2.2. IMU calibrations and cluster alignment

Gyroscope offset and magnetometer calibrations were performed for each IMU (Appendix A.1-2). IMUs were attached to rigid four-marker clusters (Figure 2) and the fixed cluster-to-IMU rotation was estimated (Appendix A.4), to generate reference orientation (*^G^q_IMU_*_,*ref*_) for tuning sensor fusion gains and assessing IMU orientation error in the Global frame.

**Figure 2.**
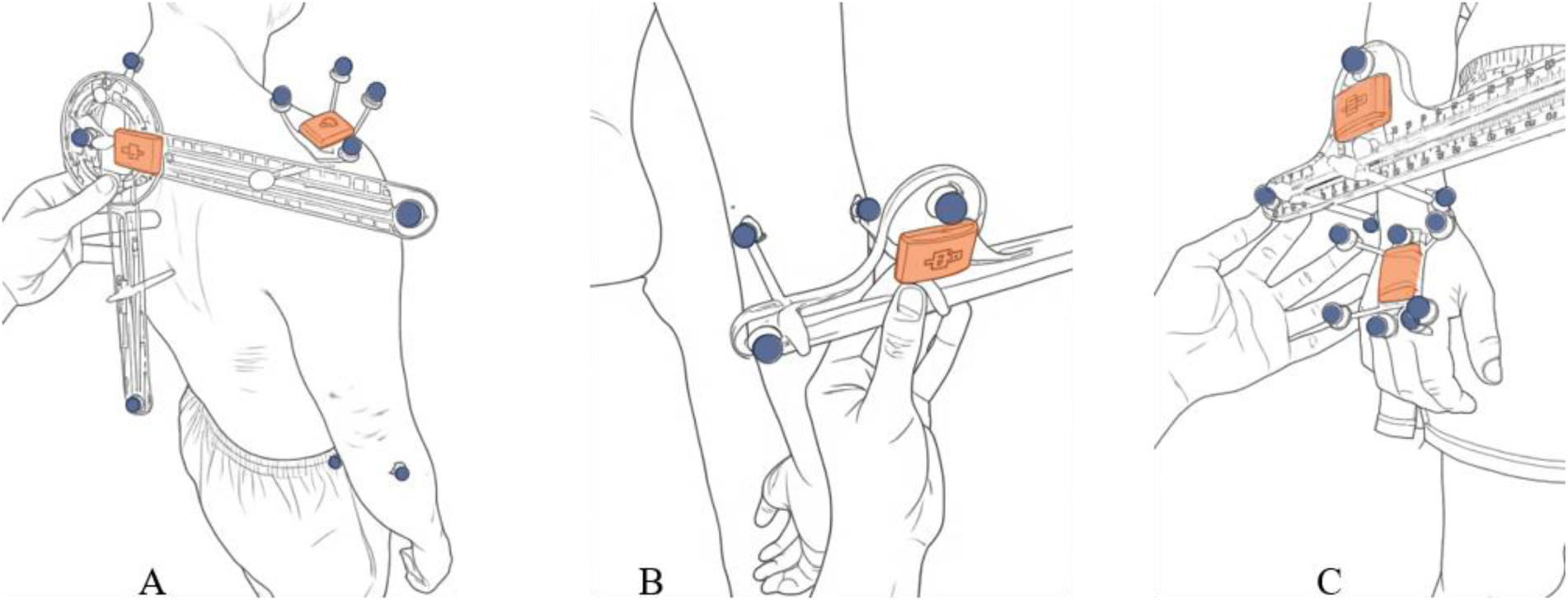
Scapula locator (A) and Calliper used for the sensor-to-segment alignment. Examples of the humerus (B) and hand (C) are shown. Orange rectangles represent IMUs and blue circles represent OMC markers.

**Figure 3.**
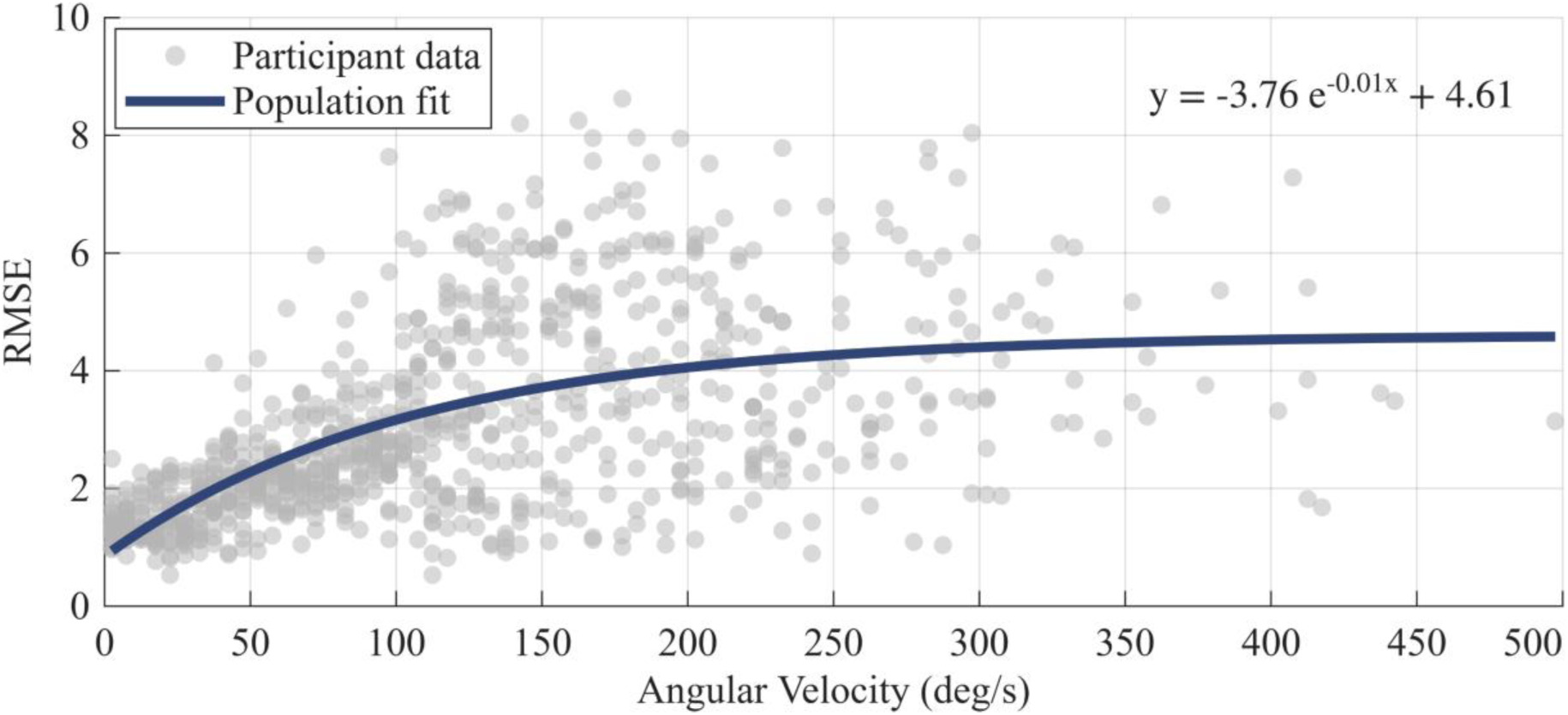
Relationship between segment angular velocity and sensor orientation RMSE. Grey circles represent individual participant data, while the blue line shows the population-level model fitted across participants (mixed-effects exponential decay model).

#### 2.2.3. Manual IMU alignment

For the manual alignment method, IMU-cluster combinations were positioned by visually aligning sensor axes with the estimated anatomical segment axes (Carnevale et al., 2019; Friesen et al., 2023; Hoglund et al., 2021; Wirth et al., 2019). They were attached to the thorax, clavicle, scapula, humerus, forearm, and hand of the dominant limb (Table 1), following placement sites intended to reduce STA (Appendix B.1).

**Table 1.**
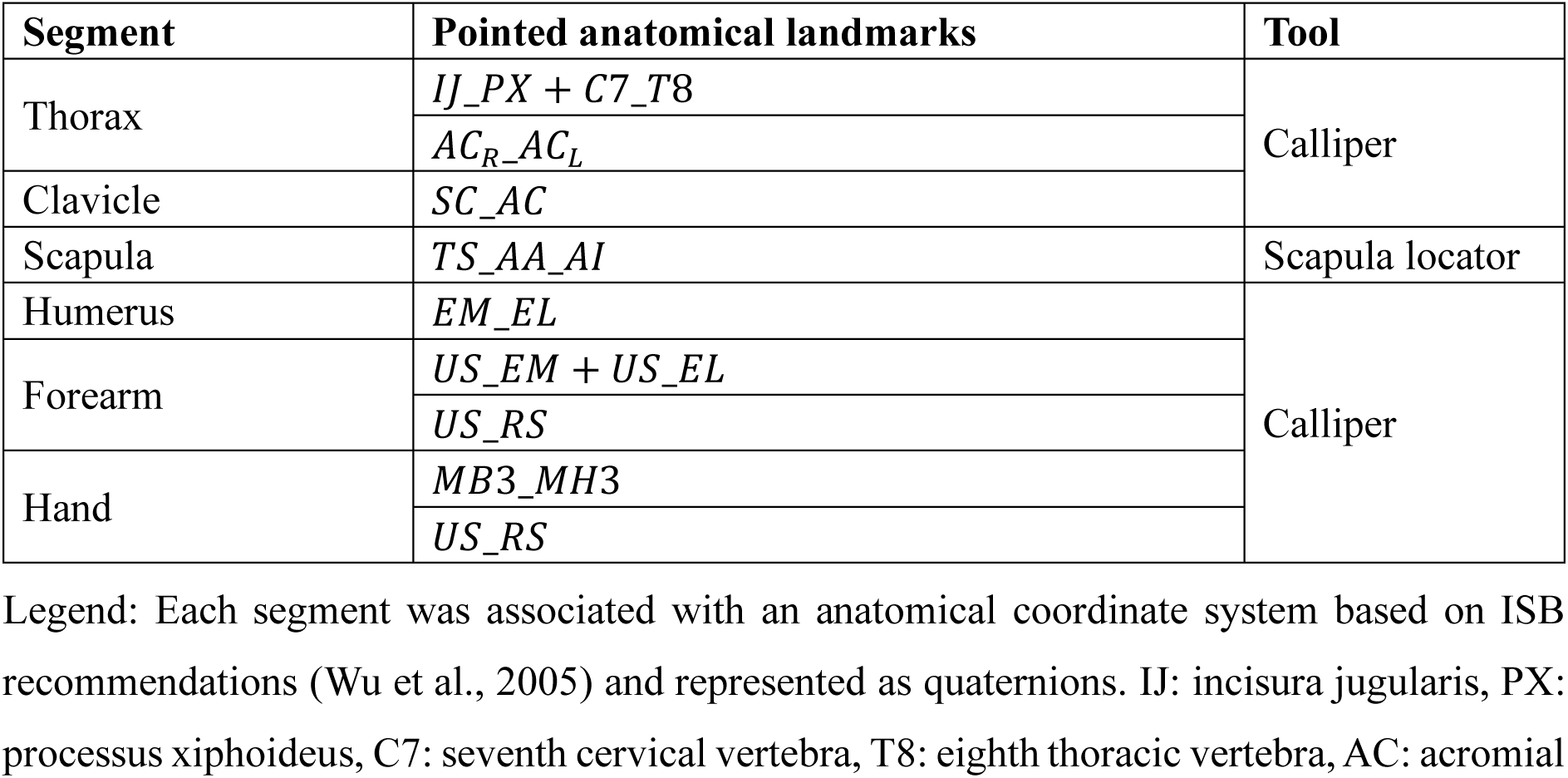
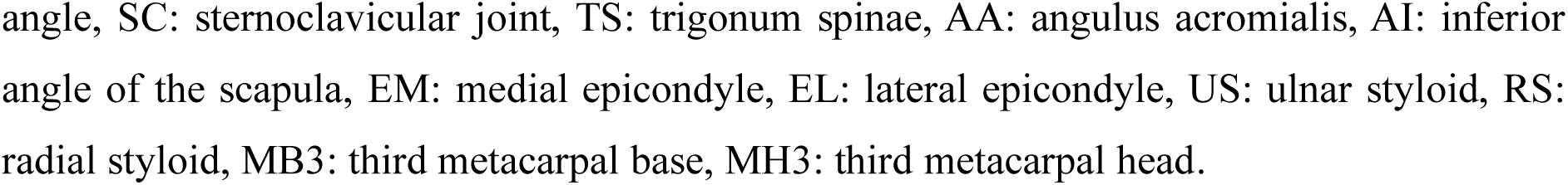
Anatomical landmarks and tools used for the IMU sensor-to-segment alignment.

#### 2.2.4. Landmark registration-based IMU alignment

Landmark registration was performed by a single experimenter while participants stood upright (Figure 2A). Custom IMU-instrumented tools were used to register anatomical landmarks according to ISB standards (Table 1): a calliper for the thorax, humerus, forearm, and wrist orientations (Figure 2B (Wu et al., 2005) and a scapula locator (Figure 2A, Appendix B.5).

Landmarks were identified at the base of the reflective markers and captured for three seconds per segment. Medial-lateral epicondyle distance was measured to estimate the epicondylar midpoint (Appendix B.3).

#### 2.2.5. Tasks

Participants first performed a static reference posture for neutral-pose alignment (upright, feet shoulder-width apart, arms resting at the sides in neutral position, head, trunk, pelvis, and limbs aligned). They then completed three repetitions of clinical and functional tasks in randomised order (Table 2), following standardised instructions and demonstration. Clinical tasks were performed through full range of motion and all tasks at self-paced speed.

**Table 2.**
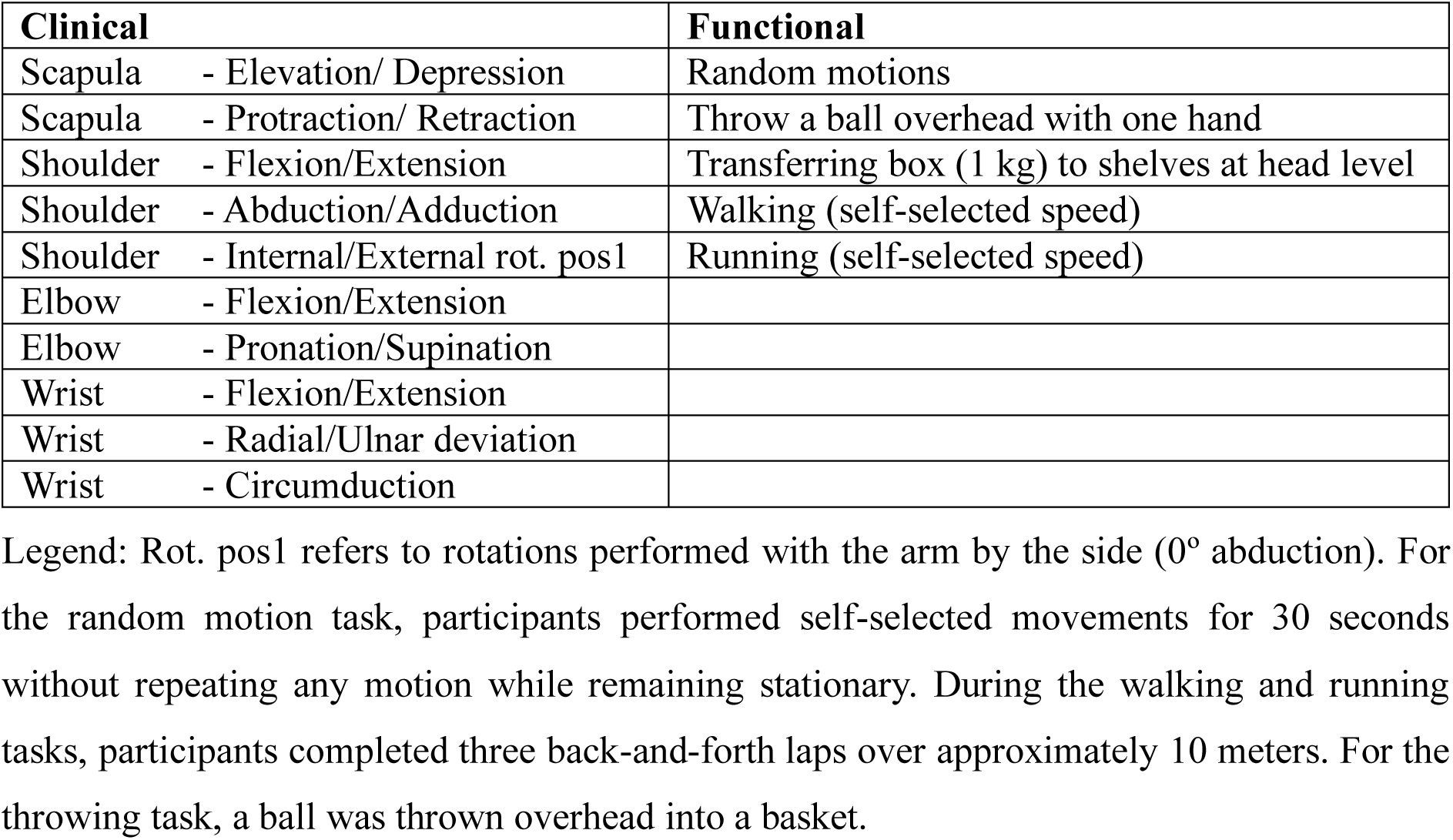
Clinical and functional tasks performed by the participants.

### 2.3. Data analysis

Data were processed offline using custom MATLAB scripts (Version R2026a, MathWorks, Natick, MA, USA). Gaps in three-dimensional marker trajectories were interpolated in Vicon Nexus using rigid body reconstruction. Marker trajectories were low-pass filtered using a second-order, bi-directional Butterworth filter with a 5 Hz cut-off frequency (Schreven et al., 2015). Unless stated otherwise, *^G^q_A_* denotes the orientation of frame *A* expressed in the global frame *G*, (·)^∗^denotes the quaternion conjugate, and ⊗ denotes quaternion multiplication.

#### 2.3.1. Aim 1: sensor fusion gain optimisation as a function of angular velocity

IMU orientation was estimated using a gradient descent filter (Riddick et al., 2023). For each IMU, sensor fusion gains were optimised offline to minimise orientation error relative to the corresponding OMC cluster-based ground truth. Optimisation (GlobalSearch.m, Matlab) was performed separately for each participant, task, and IMU-cluster unit to identify gain parameters that minimised the mean RMS orientation error across the three ZYX Cardan angle components:

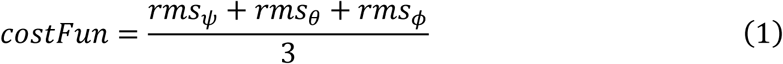

where *ψ*, *θ*, *φ* are the Cardan angle components obtained from the orientation error quaternion:

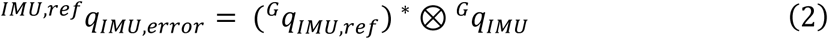

Here, *^G^q_IMU_*_,*ref*_ is the cluster-based ground truth orientation expressed in the global frame after cluster-to-IMU alignment, and *^G^q_IMU_* is the corresponding IMU orientation estimate expressed in the same frame.

To investigate how optimal gains varied with movement speed, the mean angular velocity norm was computed for each participant-task-IMU combination and modelled against optimised gains for each sensor using a mixed-effects exponential function with participant as random intercept:

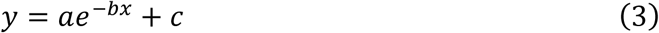

where *y* is the optimised gain, *x* the angular velocity, *a* the amplitude and direction of change, *b* the rate of change and *c* the asymptotic gain. The models were used to select the optimum sensor fusion gains based on each task’s average angular velocity norm obtained from the IMU. Final IMU orientation estimation error was then determined using Eqs. 1 & 2.

#### 2.3.2. Aim 2: Manual and landmark-based sensor-to-segment alignment accuracy

##### 2.3.2.1. Manual alignment

The global IMU orientation obtained from the fusion algorithm was used directly as the estimated global anatomical segment orientation, with the sensor axes permuted to follow the ISB segment orientation convention (see Appendix B.3 and B.4 for a detailed description of segment and joint coordinate definition).

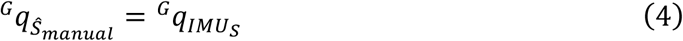

where *^G^q_Ŝ_*_manual_ is the estimated anatomical segment orientation *Ŝ*.

##### 2.3.2.2. Neutral-pose alignment

IMU orientations were corrected using a 2-s neutral reference posture:

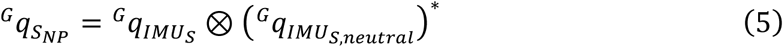

where *^G^_qIMU S,neutral_* is the mean orientation of the IMU mounted on segment *S* during the neutral-pose, and G_qSNP_ describes the neutral-pose-normalised segment orientation, not an anatomically defined segment frame.

##### 2.3.2.3. Landmark-based alignment

Anatomical axes measured with the calliper or scapula locator were expressed in the local IMU frame of the corresponding segment. Let *^G^q_IMU Tool_* be the orientation of the IMU attached to the registration tool, and *^G^q_IMUs_* the orientation of the IMU attached to segment *S*, both expressed in the global frame. For each registered anatomical axis (following ISB convention), the corresponding tool axis *e_k_*, defined by the associated anatomical landmark pair, was first expressed in the global frame:

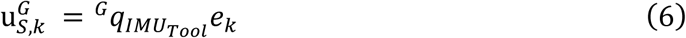

u*^G^_s,k_* was then embedded as a pure quaternion with zero scalar part and expressed in the segment *IMU* frame:

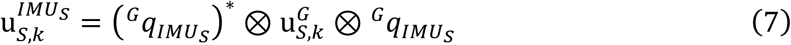

Because tool and segment IMU orientations were recorded simultaneously, each registered anatomical axis could be expressed in the corresponding segment IMU frame. Two non-collinear anatomical axes were registered for each segment (except for the clavicle which shares the thorax Z axis) and orthonormalised using the Gram-Schmidt process to define the landmark-derived anatomical segment frame expressed in the IMU frame (*^IMU^*_S_*q_Ŝ_*_landmark_, i.e. the landmark-derived sensor-to-segment transformation).

##### 2.3.2.4. Sensor-to-segment alignment error

Sensor-to-segment alignment error was calculated only for methods that assumed a fixed anatomical relationship between the IMU and the underlying segment. The neutral-pose method was excluded because it normalises IMU orientation to a reference posture and does not estimate an anatomical sensor-to-segment transformation. For each segment *S*, the OMC-derived reference sensor-to-segment transformation was calculated as:

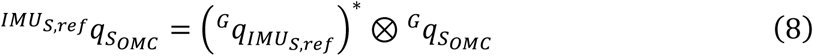

where *G_qIMUS,ref_* is the OMC-derived orientation of the segment IMU frame after cluster-to-IMU alignment, and *^G^q_SOMC_* is the OMC-derived anatomical segment orientation. The sensor-to-segment error was then calculated as:

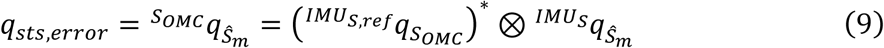

Where *m* ∈ *M_anat_* = {*manual*, *landmark*}. The error quaternion *q_sts_*_,*error*_ was decomposed using a ZYX sequence, and the RMS of the three decomposed axes was calculated for each alignment method, segment, movement, and participant.

#### 2.3.3. Aim 3: Accuracy of the manual, neutral-pose and landmark-based registration methods for joint angle estimation

For each joint *J* of each alignment method *m* ∈ {*OMC*, *manual*, *landmark*, *neutral pose*}, the estimated joint orientation was calculated as the orientation of the distal segment *D* relative to the proximal segment *P*.

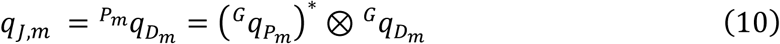

Joint angle error for IMU-based alignment method *m* ∈ {*manual*, *landmark*, *neutral pose*} was then calculated as:

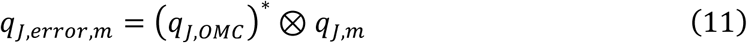

The resulting error quaternion *q_J_*_,*error*,*m*_ was decomposed using a ZYX sequence, and RMSE was calculated across axes, joints, movements, and alignment methods.

#### 2.3.4. Statistics

All statistical analyses were performed in RStudio (Version 2026.01.1+403, Posit Software, Boston, MA, USA), with statistical significance set at p≤0.05. Error outcomes were the RMSE of decomposed quaternion error components, averaged across axes unless otherwise stated, and reported as mean with 95% confidence intervals (CI).

Sensor-to-segment and joint angle errors were each analysed using linear mixed-effects models (nlme package), with alignment method, segment or joint, movement, and their interactions as fixed effects, participant as random intercept, and a segment-or joint-specific residual variance structure (varIdent) to account for heteroscedasticity. Where interactions were significant, estimated marginal means (EMM, emmeans package) were calculated, with Bonferroni-adjusted post-hoc comparisons used to compare alignment methods within each segment-or joint-movement combination.

Bland-Altman analyses were performed on each Cardan angle component to characterise the difference between IMU-and OMC-derived joint angles (i.e. bias), with limits of agreement (LoA) calculated across range of motion. Mixed-effects models were fitted to the Bland–Altman data to quantify proportional bias.

## 3. Results

### 3.1. Aim 1: Fusion parameters optimisation and sensor orientation errors

Optimal fusion gains varied with angular velocity and were well described by exponential models for all sensors (adjusted *R*^2^ ≥ 0.95, Table 3, Appendix C.1). Gyroscope gain increased with angular velocity toward an asymptote of *c* = 0.97, while accelerometer and magnetometer gains decreased towards asymptotes of *c* = 0.258 and *c* = 0.213, respectively. This pattern indicates that faster movements were weighted more toward gyroscope integration and less toward accelerometer-and magnetometer-based correction.

**Table 3.** Fitted parameters of the exponential model *y* = *ae*^−*bx*^ + *c* describing the relationship between angular velocity and fusion gain for each sensor type.

| Sensor | Exponential model parameters | | | $R^2$ |
| --- | --- | --- | --- | --- |
| | $a$ [95% CI] | $b$ [95% CI] | $c$ [95% CI] | |
| Gyroscope | -0.48 [-0.62 - -0.39] | 0.13 [0.11 - 0.16] | 0.97 [0.96 - 0.98] | 0.99 |
| Accelerometer | 0.43 [0.360 - 0.49] | 0.03 [0.02 - 0.04] | 0.26 [0.22 - 0.29] | 0.95 |
| Magnetometer | 0.46 [0.39 - 0.53] | 0.02 [0.02 - 0.03] | 0.21 [0.17 - 0.25] | 0.98 |

Legend: Parameters are reported with 95% CI. Adjusted $R^2$ indicates model goodness of fit.

### 3.2. Aim 2: Sensor-to-segment alignment error

Sensor-to-segment RMSE differed between alignment methods across all segments (Figure 4), with landmark-based alignment significantly lower (2.0° to 5.4°) than manual alignment (6.7° to 19.4°) for every segment (F=9.40, Landmark-Manual=-9.3° CI=[-9.4,-9.2], all p<0.0001). Between-method differences were largest at the scapula and humerus and smallest at the hand, where a few movement-specific comparisons were not significant.

**Figure 4.**
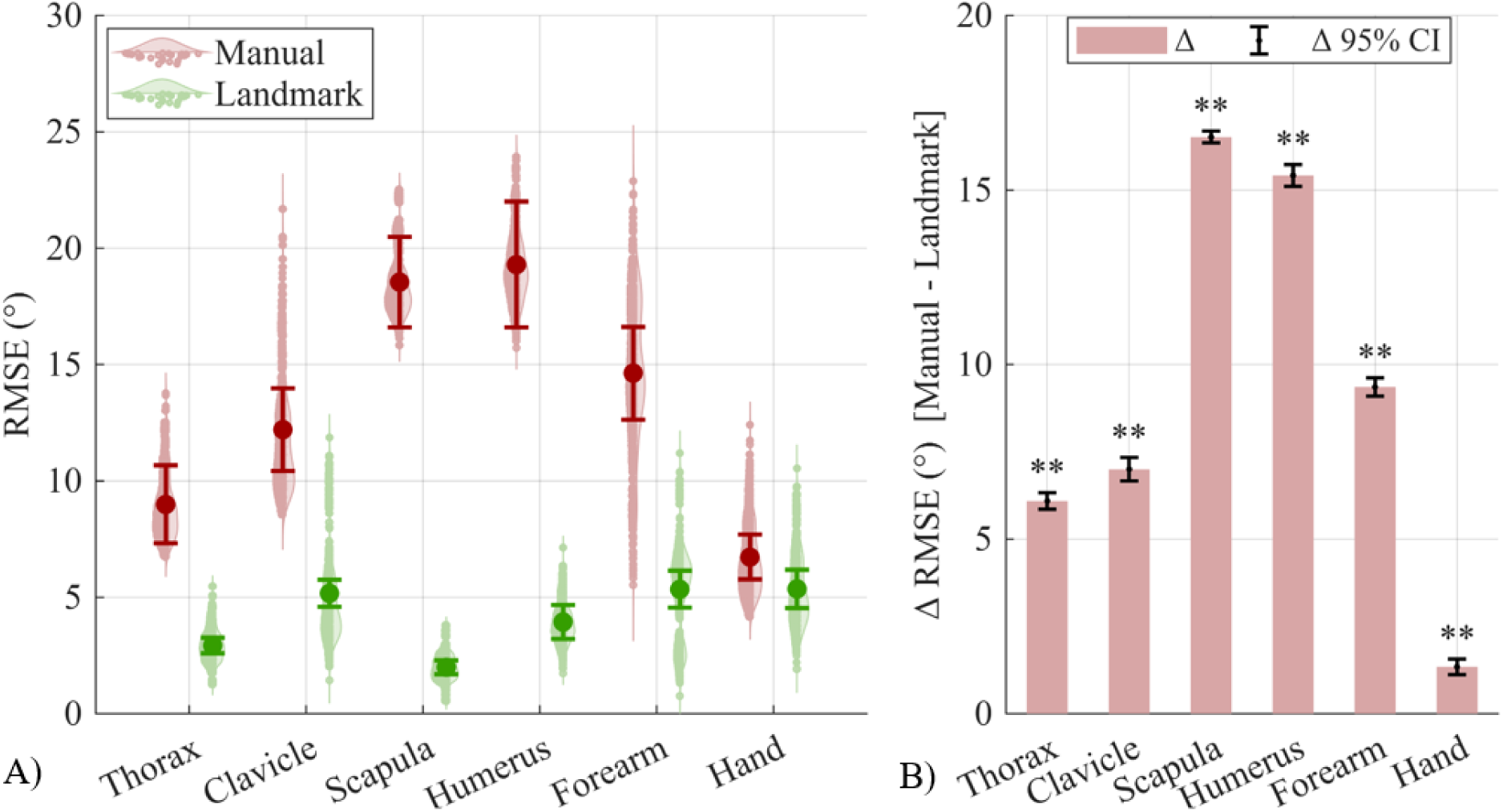
Sensor-to-segment alignment error across all tasks for each segment and alignment method. A) Violin plots represent conditional fitted values derived from linear mixed-effects models. Solid circles with error bars denote EMM and 95% CI. B) Between-method difference in RMSE per segment. Significance stars denote the outcome of pairwise comparison between methods: ** p<0.01.

### 3.3. Aim 3: Joint angle errors

#### 3.3.1. Accuracy

Landmark-based alignment yielded the lowest joint angle RMSE overall, from 3.7° 95% CI [3.2-4.2°] at the scapulothoracic joint to 7.8° [7.0-8.5°] at the elbow (Figure 5), significantly lower than manual and neutral-pose alignment at every joint (Method×Joint×Movement F=20.9, Landmark-Manual=-13.4, Landmark-Neutral=-17.5: all, p<0.0001). Manual alignment errors were highest at the acromioclavicular (24.1° [23.5-24.6°]), elbow (24.2° [23.5-25.0°]), and glenohumeral joints (22.1° [21.4-22.8°]), and lowest at the sternoclavicular joint (8.3° [7.7-8.9°]). Neutral-pose alignment produced the highest errors overall, particularly at the elbow (35.5° [34.8-36.3°]) and humerothoracic joint (28.0° [27.3-28.7°]). Manual and neutral-pose mean errors were small at the scapulothoracic joint (Δ=0.4°, p=0.042) and similar at the glenohumeral (Δ=0.6°, p=0.250) and wrist (Δ=-0.3°, p=0.457).

**Figure 5.**
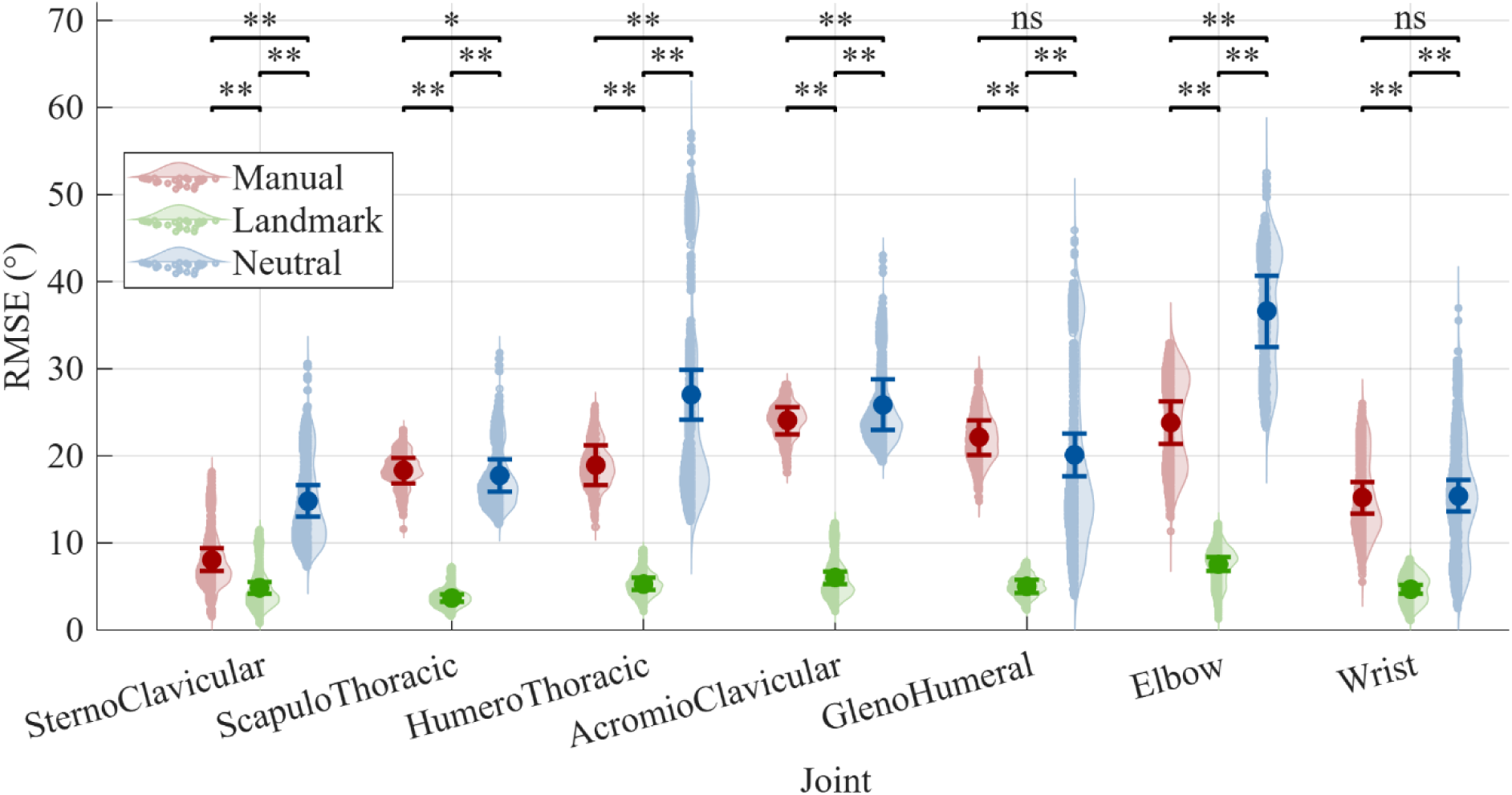
Joint-specific RMSE of the alignment methods across all movements. Violin plots represent conditional fitted values derived from linear mixed-effects models. Circles with error bars denote EMM and 95% CI.

#### 3.3.2. Landmark-based alignment agreement

Figure 6 characterises agreement between landmark-based and OMC joint angles using Bland-Altman analysis. Mean bias (i.e. OMC-IMU) was generally small and centred near zero, although several joints showed proportional bias across joint angle magnitude.

**Figure 6.**
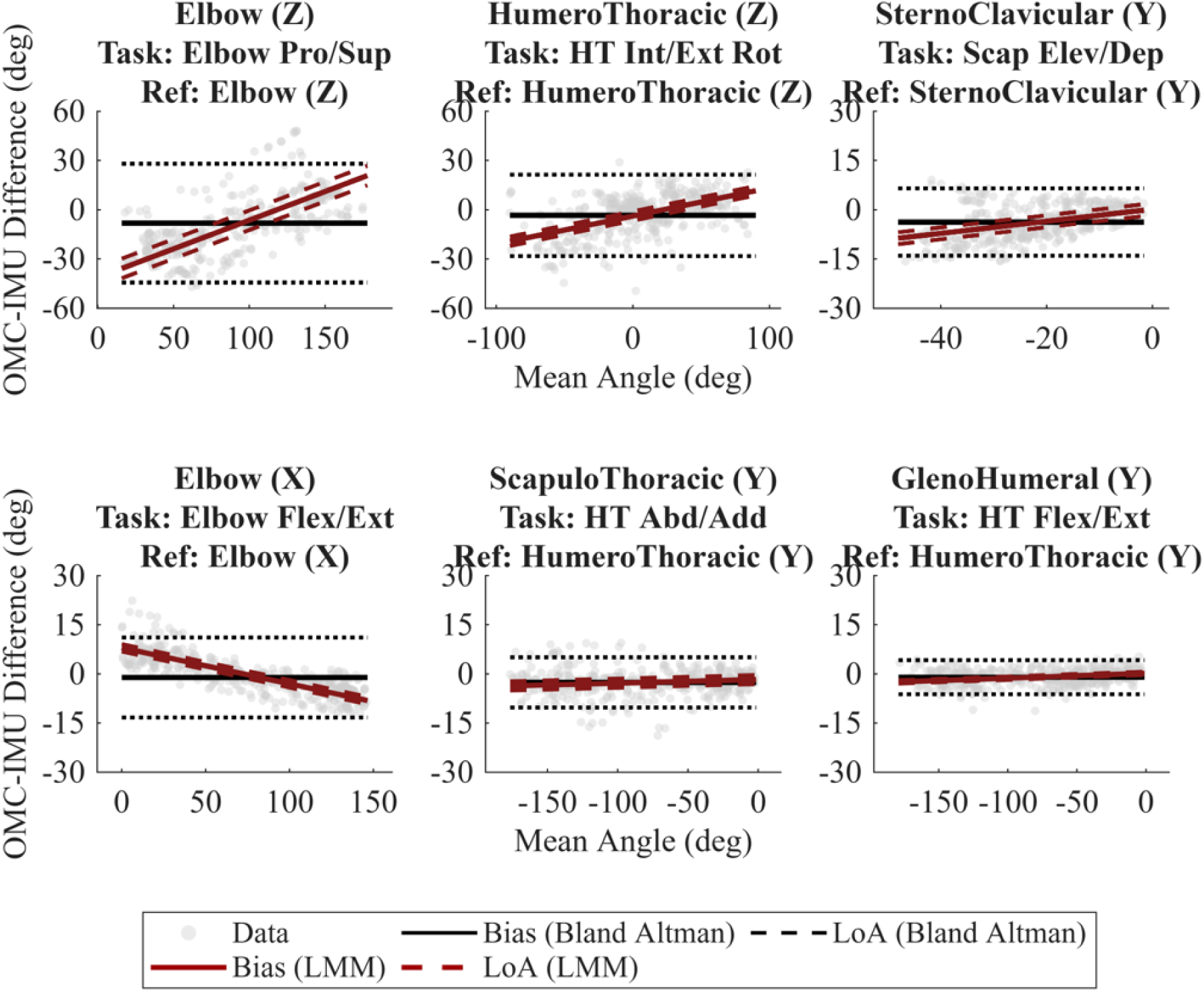
Bland-Altman comparison of representative joint angles between the landmark-based alignment and the OMC reference system. Title lines indicate in order: joint of the measured angle error, the task performed, and the reference joint/axis against which it was plotted. Black solid and black dashed lines show Bland-Altman mean bias and limits of agreement; red line and red dashed lines show linear mixed model-estimated proportional bias and its 95% CI. Positive bias indicates IMU underestimation relative to OMC. Full Bland-Altman analyses for all Joint×Movement combinations are shown in Appendix C.2 and C.3. Flex/Ext: Flexion/Extension, Abd/Add: abduction/adduction, Int/Ext Rot: internal/external rotation, Scap Ele/Dep: scapular elevation/depression.

Proportional bias was largest for elbow pronation/supination (β=0.349°/° CI=[0.347, 0.351], intercept=-41.3° CI=[-47.3,-35.3]), humerothoracic internal/external rotation (β=0.171°/° [0.169, 0.173], intercept=-3.8° [-6.6,-1.0]) and sternoclavicular scapular elevation/depression (β=0.184°/° [0.180, 0.187], intercept=0.2° [-1.6, 2.0]), reflecting greater IMU underestimation with increasing angle. Elbow flexion/extension showed the opposite pattern (β=-0.110°/° [-0.111,-0.109], intercept=8.0° [6.8, 9.1]). Bias was smaller for scapulothoracic abduction/adduction (β=0.011°/° [0.011, 0.012], intercept=-1.7° [-3.1,-0.3]) and glenohumeral flexion/extension (β=0.015°/° [0.014, 0.016], intercept=0.2° [-0.3, 0.7]).

Bland-Altman bias and LoA were widest for elbow pronation/supination (-8.2° [-44.3, 28.0]) and humerothoracic internal/external rotation (-3.4° [-28.2, 21.4]), and narrowest for scapulothoracic elevation/depression (-0.0° [-4.5, 4.4]) and glenohumeral abduction/adduction (-0.2° [-4.8, 4.4]).

#### 3.3.3. Landmark-based alignment joint/task performance

Landmark-based alignment joint angle error varied across tasks (Joint×Movement interaction F=27.5, p<0.0001, Figure 7). Complete EMMs are provided in Appendix C.4-C.6. Dominant patterns and largest errors within clinical and functional tasks are summarised below.

**Figure 7.**
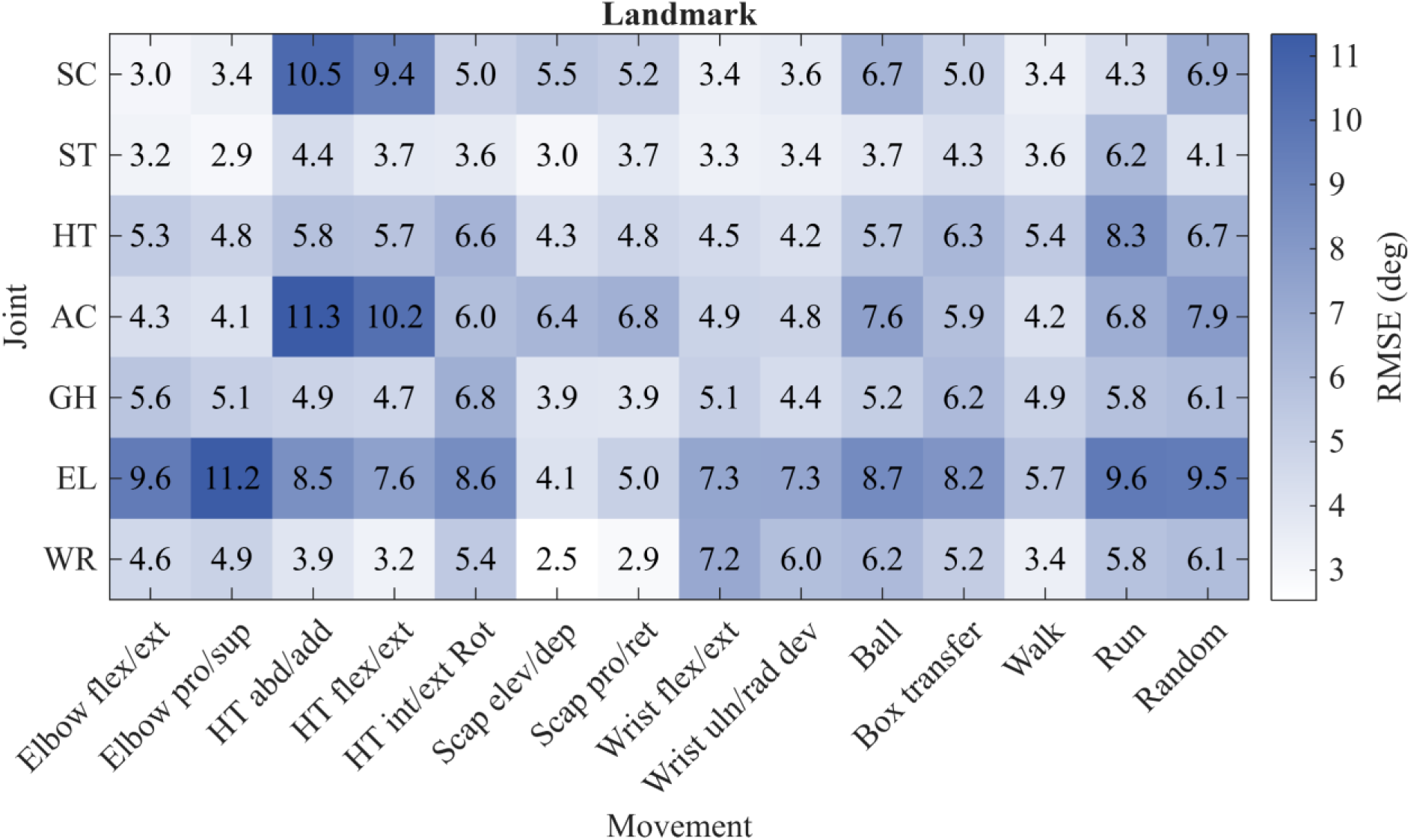
Heatmap representation of the linear mixed model-derived EMMs across each Joint×Movement combination for the landmark-based alignment method. Flex/ext: flexion/extension, Pro/sup: protraction/supination, Abd/add: abduction/adduction, Int/ext rot: internal/external rotation, Scap: scapular, Elev/dep: elevation/depression, Pro/ret: protraction/retraction, Uln/rad dev: ulnar/radial deviation, Circ: circumduction, SC: sternoclavicular, ST: scapulothoracic, AC: acromioclavicular, GH: Glenohumeral, HT: humerothoracic, EL: elbow, WR: wrist.

Clinical task errors were generally highest at the joint most engaged in the movement. Humerothoracic abduction/adduction and flexion/extension produced the largest errors at the acromioclavicular (11.3°, 10.2°) and sternoclavicular joints (10.5°, 9.4°), with all other joints below 8.5°. Scapula elevation/depression and protraction/retraction showed a similar pattern, highest at the acromioclavicular (6.4°, 6.6°) and sternoclavicular (5.5°, 5.1°) joints, with low scapulothoracic error (<3.6°). Elbow errors peaked during elbow flexion/extension (9.6°) and pronation/supination (11.2°), and wrist tasks showed errors across both wrist and elbow (6° to 8.2°).

Compared with clinical tasks, errors during functional tasks were more evenly distributed across joints. Running produced the largest proximal errors, at the humerothoracic (8.29°) and scapulothoracic (6.3°) joints. Walking produced lower, more uniform errors across joints (3.4°-5.7°). During throwing, box transfer, and random movement, the elbow showed the highest errors (8.7°-9.5°, respectively), followed by the acromioclavicular joint (7.6°-7.9°), while the scapulothoracic joint showed the lowest ones (<4.2°). The sternoclavicular joint showed moderate but persistent error across most functional tasks (3.4°-6.9°).

## 4. Discussion

This study developed and tested the accuracy of a multi-segment IMU framework for estimating anatomically meaningful upper-limb joint angles by evaluating three sequential error sources: sensor orientation estimation, sensor-to-segment alignment, and final joint angle accuracy. Sensor orientation errors were low after velocity-dependent fusion-gain optimisation, indicating minimal IMU orientation estimation contribution to final joint error. In contrast, joint angle accuracy depended strongly on alignment method, joint, and movement task. Compared with landmark-based registration, manual and neutral-pose alignment exhibited larger errors or joint angles less directly linked to anatomical coordinate systems. Therefore, alignment affects joint-angle interpretability.

Sensor fusion optimisation produced low orientation errors across segments, with mean error remaining below 3°. Although gains varied systematically with angular velocity, this mainly confirms that sensor orientation estimation was sufficiently accurate to support the subsequent validation steps. The larger and more clinically relevant errors emerged later, particularly during sensor-to-segment alignment and joint-angle calculation.

The sensor-to-segment alignment results highlight that the method is central to the anatomical and therefore clinical interpretability of IMU-derived upper-limb kinematics, consistent with prior work showing that the alignment method affects upper-limb IMU-derived joint angle accuracy (Bouvier et al., 2015; Fang et al., 2023). Manual alignment relies on the operator visually approximating anatomical axes from sensor placement, which is vulnerable to palpation error, body morphology, STA, and the complex shape of shoulder-girdle segments (Bonfiglio et al., 2024). This likely explains why manual alignment errors propagate into substantially larger joint-angle errors compared to the landmark-based alignment. In contrast, landmark-based registration explicitly defines segment coordinate systems from anatomical landmarks, directly linking sensor orientation and anatomical joint definitions (Picerno et al., 2019; Wu et al., 2005). This distinction matters clinically: only anatomically defined joint angles are required for absolute anatomical interpretation and provide a more appropriate basis for between-participant and longitudinal comparison (Fang et al., 2023; Wu et al., 2005). The present study builds on previous bony-landmark approaches by applying the method across the full upper-limb chain during both clinical and functional movements (Friesen et al., 2023; Picerno et al., 2019). Neutral-pose alignment removes initial orientation offsets and may be useful in time-constrained settings, but does not define the sensor orientation relative to the underlying anatomical segment (Bouvier et al., 2015; Fang et al., 2023). Therefore, even when neutral-pose alignment produces acceptable joint angle errors, its angles are less suitable for absolute anatomical interpretation or cross-sectional/longitudinal comparison.

Joint angle errors were not uniform across the upper limb, but followed a structured joint-and task-dependent pattern, supporting previous observations that IMU-derived upper-limb kinematics are sensitive to both alignment approach and movement context (Bouvier et al., 2015; Fang et al., 2023). During isolated clinical tasks, errors tended to be greatest at the joint being moved or at mechanically coupled joints. This likely reflects the anatomical and measurement complexity of the shoulder girdle, where limited sensor placement area, soft tissue displacement, and mechanical coupling between adjacent segments can amplify the effect of small segment-orientation errors on sternoclavicular and acromioclavicular joint angles (Begon et al., 2018; Duprey et al., 2017). In contrast, functional tasks produced a more distributed error pattern across the kinematic chain, consistent with their multi-planar and multi-joint nature. Running was the clearest example, with elevated humerothoracic and scapulothoracic errors likely reflecting fast cyclical motion, whole-body acceleration, and greater dynamic soft tissue movement. Walking produced a more uniform and lower error profile, suggesting that the proposed framework is better suited to controlled clinical movements and lower-speed functional tasks than to fast dynamic activities. Bland-Altman analysis further showed that agreement between IMU-derived and OMC-derived angles was not constant across the full angular range for all joints. These angle-dependent discrepancies likely reflect STA during large-amplitude movements and are important because they indicate that errors may be small near neutral but increase, or change direction, at larger joint excursions. This should be considered when interpreting end-range motion, particularly in clinical assessments where deficits or compensatory strategies may be most apparent at the limits of available movement.

Although this staged validation framework enables the localisation of orientation error sources, several limitations should be considered. First, sensor-fusion parameters were optimised in a controlled laboratory environment and may not generalise to real-world settings where magnetic disturbances can degrade heading estimation, particularly if magnetometer weighting or calibration is inadequate. Second, the filter was initialised from an identity quaternion rather than the true sensor orientation. Although likely brief, this transient may affect short-duration tasks unless a stabilisation period precedes movement. Third, landmark-based registration depends on accurate palpation and tool placement, introducing operator dependence, particularly at the scapula and clavicle. The approach requires greater anatomical expertise and offline processing than manual or neutral-pose alignment, which may limit routine clinical use unless alignment is streamlined. Fourth, the OMC reference used skin-mounted markers and clusters and therefore does not provide direct bone-motion accuracy. Both systems are susceptible to STA, and some errors may be shared. In addition, landmarks were registered at the bases of OMC markers, and scapular alignment used the same locator-derived anatomical definition, reducing independence between methods and potentially favouring agreement. Accordingly, the reported errors reflect agreement with the OMC implementation rather than absolute skeletal accuracy. This is relevant for the scapula, where low between-system disagreement should not be interpreted as equivalent skeletal accuracy. Fifth, validation was performed in participants with typical shoulder function. Accuracy may differ in clinical populations, because restricted range of motion, muscle atrophy, altered movement strategies, or post-surgical anatomical changes may hinder landmark identification and sensor placement. Finally, repeatability and inter-operator agreement were not evaluated, so sensitivity to examiner differences and repeated testing sessions remains unknown.

## 5. Conclusion

This study validated a multi-segment IMU framework for estimating anatomically meaningful upper-limb joint angles. Sensor orientation errors were low after velocity-dependent fusion-gain optimisation, but final joint-angle accuracy depended strongly on alignment method, joint, and task performed. Landmark-based alignment showed the most accurate and anatomically interpretable joint angles, providing the most promising basis for further clinical evaluation. These findings show that IMU-based upper limb assessment has clinical potential, provided accuracy is interpreted in relation to the specific alignment method, joint, and movement context.

## Author contributions

### Arthur Fabre

Conceptualization, Data curation, Formal analysis, Investigation, Methodology, Project administration, Resources, Software, Supervision, Visualization, Writing – original draft, Writing – review and editing.

### Graham Kerr

Conceptualization, Formal analysis, Funding acquisition, Methodology, Project administration, Resources, Supervision, Writing – review and editing.

### Peter Pivonka

Funding acquisition, Supervision, Writing – review and editing.

### Wolbert van den Hoorn

Conceptualization, Formal analysis, Methodology, Project administration, Resources, Software, Supervision, Visualisation, Writing – original draft, Writing – review and editing.

## Supporting information

Appendixes

## Acknowledgments

The authors gratefully acknowledge the support of the (ARC ITTC-JB, IC190100020).

## References

Begon, M., Andersen, M. S., & Dumas, R. (2018). Multibody kinematics optimization for the estimation of upper and lower limb human joint kinematics: a systematized methodological review. J Biomech Eng, 140(3). 10.1115/1.4038741

Bonfiglio, A., Tacconi, D., Bongers, R. M., & Farella, E. (2024). Effects of IMU sensor-to-segment calibration on clinical 3D elbow joint angles estimation. Front Bioeng Biotechnol, 12, 1385750. 10.3389/fbioe.2024.1385750

Bouvier, B., Duprey, S., Claudon, L., Dumas, R., & Savescu, A. (2015). Upper limb kinematics using inertial and magnetic sensors: comparison of sensor-to-segment calibrations. Sensors, 15(8), 18813–18833. 10.3390/s150818813

Carnevale, A., Longo, U. G., Schena, E., Massaroni, C., Lo Presti, D., Berton, A., Candela, V., & Denaro, V. (2019). Wearable systems for shoulder kinematics assessment: a systematic review. BMC Musculoskeletal Disorders, 20(1), 546. 10.1186/s12891-019-2930-4

Crabolu, M., Pani, D., Raffo, L., Conti, M., Crivelli, P., & Cereatti, A. (2017). In vivo estimation of the shoulder joint center of rotation using magneto-inertial sensors: MRI-based accuracy and repeatability assessment. Biomed Eng Online, 16(1), 34. 10.1186/s12938-017-0324-0

Donnelly, T. D., Ashwin, S., Macfarlane, R. J., & Waseem, M. (2013). Clinical assessment of the shoulder. Open Orthop J, 7, 310–315. 10.2174/1874325001307010310

Duprey, S., Naaim, A., Moissenet, F., Begon, M., & Cheze, L. (2017). Kinematic models of the upper limb joints for multibody kinematics optimisation: An overview. J Biomech, 62, 87–94. 10.1016/j.jbiomech.2016.12.005

Ehrig, R. M., Taylor, W. R., Duda, G. N., & Heller, M. O. (2006). A survey of formal methods for determining the centre of rotation of ball joints. J Biomech, 39(15), 2798–2809. 10.1016/j.jbiomech.2005.10.002

Fang, Z., Woodford, S., Senanayake, D., & Ackland, D. (2023). Conversion of Upper-Limb Inertial Measurement Unit Data to Joint Angles: A Systematic Review. Sensors (Basel), 23(14). 10.3390/s23146535

Friesen, K. B., Sigurdson, A., & Lang, A. E. (2023). Comparison of scapular kinematics from optical motion capture and inertial measurement units during a work-related and functional task protocol. Med Biol Eng Comput, 61(6), 1521–1531. 10.1007/s11517-023-02794-2

Hayes, K., Walton, J. R., Szomor, Z. R., & Murrell, G. A. (2001). Reliability of five methods for assessing shoulder range of motion. Aust J Physiother, 47(4), 289–294. 10.1016/s0004-9514(14)60274-9

Hindle, B. R., Keogh, J. W. L., & Lorimer, A. V. (2021). Inertial-based human motion capture: a technical summary of current processing methodologies for spatiotemporal and kinematic measures. Applied Bionics and Biomechanics, 2021, 6628320. 10.1155/2021/6628320

Hoglund, G., Grip, H., & Ohberg, F. (2021). The importance of inertial measurement unit placement in assessing upper limb motion. Medical Engineering & Physics, 92, 1–9. 10.1016/j.medengphy.2021.03.010

Madgwick, S. O., Harrison, A. J., & Vaidyanathan, A. (2011). Estimation of IMU and MARG orientation using a gradient descent algorithm. IEEE Int Conf Rehabil Robot, 2011, 5975346. 10.1109/ICORR.2011.5975346

Picerno, P., Caliandro, P., Iacovelli, C., Simbolotti, C., Crabolu, M., Pani, D., Vannozzi, G., Reale, G., Rossini, P. M., Padua, L., & Cereatti, A. (2019). Upper limb joint kinematics using wearable magnetic and inertial measurement units: an anatomical calibration procedure based on bony landmark identification. Scientific Reports, 9(1), 14449. 10.1038/s41598-019-50759-z

Riddick, R., Smits, E., Faber, G., Shearwin, C., Hodges, P., & van den Hoorn, W. (2023). Estimation of human spine orientation with inertial measurement units (IMU) at low sampling rate: How low can we go? J Biomech, 157, 111726. 10.1016/j.jbiomech.2023.111726

Schreven, S., Beek, P. J., & Smeets, J. B. (2015). Optimising filtering parameters for a 3D motion analysis system. J Electromyogr Kinesiol, 25(5), 808–814. 10.1016/j.jelekin.2015.06.004

Stefana, E., Marciano, F., Rossi, D., Cocca, P., & Tomasoni, G. (2021). Wearable devices for ergonomics: a systematic literature review. Sensors, 21(3). 10.3390/s21030777

Wirth, M. A., Fischer, G., Verdu, J., Reissner, L., Balocco, S., & Calcagni, M. (2019). Comparison of a new inertial sensor based system with an optoelectronic motion capture system for motion analysis of healthy human wrist joints. Sensors, 19(23). 10.3390/s19235297

Wu, G., van der Helm, F. C., Veeger, H. E., Makhsous, M., Van Roy, P., Anglin, C., Nagels, J., Karduna, A. R., McQuade, K., Wang, X., Werner, F. W., Buchholz, B., & Biomechanics, I. S. o. (2005). International Society of Biomechanics recommendation on definitions of joint coordinate systems of various joints for the reporting of human joint motion-Part II: shoulder, elbow, wrist and hand. Journal of biomechanics, 38(5), 981–992. 10.1016/j.jbiomech.2004.05.042

Zhu, K., Li, J., Li, D., Fan, B., & Shull, P. B. (2023). IMU shoulder angle estimation: Effects of sensor-to-segment misalignment and sensor orientation error. IEEE Trans Neural Syst Rehabil Eng, 31, 4481–4491. 10.1109/TNSRE.2023.3331238

