## Appendixes for "IMU-based upper limb kinematics validation: sensor fusion optimisation, sensor-to-segment alignment, and joint angle accuracy"

The appendix is structured into three sections.

- Section A provides an overview of device and reference frame calibration procedures, including gyroscope bias correction, magnetometer calibration for soft- and hard-iron distortions, and alignment of the global optoelectronic motion capture (OMC) reference frame with gravity and magnetic north, and alignment of each local OMC marker-cluster frame and inertial measurement unit (IMU) reference frames at both global and local levels.
- Section B includes a tabulation of sensor placement sites, manual alignment procedures for each body segment, estimation of the glenohumeral joint rotation centre, definitions of coordinate systems and joint rotation sequences used across four alignment methods (OMC, Neutral-pose, Manual, and Landmark registration), and presents the scapula orientation estimation pipeline.
- Section C summarises the analysis results, including the exponential mixed-effects model of sensor fusion gain as a function of angular velocity, Bland-Altman plots and agreement parameters for the Landmark alignment method, and heatmap visualisations with comprehensive tables of estimated marginal means of joint angle root mean square error across all joint–movement combinations for all alignment methods.

### A. Device and system calibrations

Upper limb movements were recorded in a volume ($216m^3$) surrounded by a 12-camera OMC system mounted at a height of 3.05 m. OMC and IMU data were collected (Vicon Nexus, v.2.15) and time-synchronised via Vicon Beacon hardware (version 1, firmware v.810).

Both the accuracy of sensor orientation estimation and its comparison with OMC-derived orientation are influenced by several factors, including:

- Gyroscope bias integration, a characteristic inherent to the sensor during manufacturing, can result in an accumulating low-frequency drift in orientation estimation.
- Ferromagnetic disturbances in the surrounding environment, causing the signal to be corrupted by high-frequency noise, result in error in the heading correction.
- Misalignment between the global OMC Z-axis and the gravity axis, affecting the orientation of the XY-plane.
- Local misalignment between the cluster and sensor-derived orientations.
- Misalignment between the global OMC X-axis and the local magnetic north direction, determined by the sensors' magnetometer.

To minimise errors associated with each factor and to facilitate comparison between sensor and OMC orientations, the following sections outline the subsequent corresponding adjustments.

#### A.1. Gyroscope bias

For each sensor, a static 20-second trial is recorded before each data acquisition, and the mean value along each axis is subtracted from the final gyroscope readings (Figure 1).

Figure 1. Tri-axial gyroscope output recorded during a static calibration trial

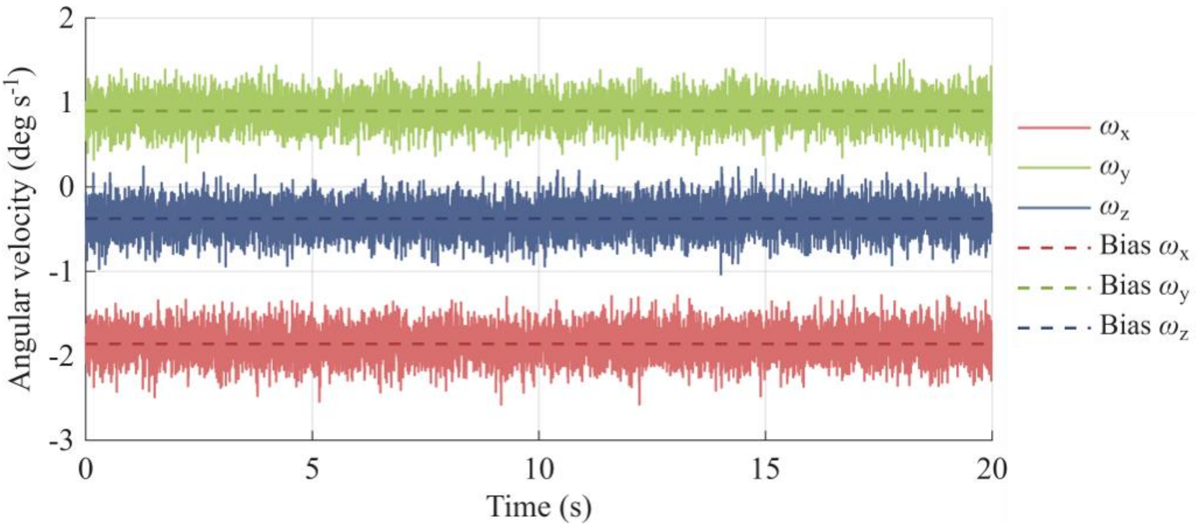


Legend: Angular velocity ($\omega$) recorded from a representative inertial measurement unit (IMU) gyroscope (x, y, z axes) during a 20-s static trial performed before each data-collection session. The non-zero offset from 0°/s along each axis reflects the sensor-specific gyroscope bias, the mean of which was subtracted from all subsequent gyroscope recordings within that session

#### A.2. Ferromagnetic disturbances

The ideal measurements from a magnetometer would form a perfect sphere centred at the origin in a 3-D rotation scenario, assuming an unperturbed magnetic field. However, distortions from the sensor circuit board and surrounding environment can cause deviations. Two main effects are observed: the soft-iron effect, which distorts the spherical shape into an ellipsoid and introduces tilt, often caused by external disturbances influencing the magnetic field; and the hard-iron effect, which results in an offset of the ellipsoid centre from the origin, typically caused by materials with a constant, additive magnetic field contribution to the Earth's magnetic field. The magnetometer calibration process involves a 120-second capture before each data collection, during which each sensor is rotated along the $XYZ$ axis for thirty seconds each, followed by random rotations. Then, the MATLAB function "magcal" is used to obtain calibration coefficients and correct for any disturbances (Figure 2).

Figure 2. Magnetometer calibration for soft- and hard-iron distortion.

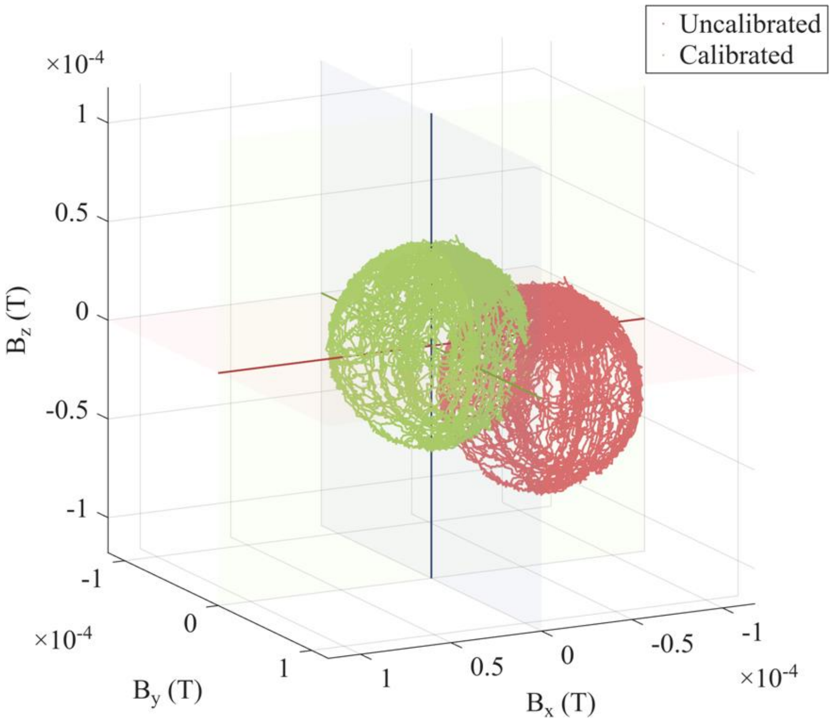


Legend: Raw (red) and calibrated (green) magnetometer measurements from a representative IMU, plotted in three-dimensional magnetic flux space (T).

#### A.3. Misalignment between OMC z-axis and gravity

Although the calibration wand was levelled as carefully as possible before establishing the OMC volume origin, residual misalignment between the OMC-defined Z-axis and true vertical (gravity) could not be entirely excluded. To correct for this, a plumb-line pendulum threaded through two reflective markers was used to establish the gravitational vector in the OMC volume (Figure 3). This procedure was performed before each data collection session, and the mean three-dimensional position of each marker over 2 minutes was used to define a unit vector ${}^{G}v_{g_z}$ representing true vertical. This vector served as the first basis vector in a Gram-Schmidt orthonormalisation process, together with the OMC reference frame axes ${}^{G}v_{OMC_x}$ and ${}^{G}v_{OMC_y}$, to construct a correction quaternion that aligned the OMC global $(G)$ Z-axis with the estimated gravity vector (Figure 3).

$$ {}^{G}v_{g_x} = \frac{\left({}^{G}v_{OMC_y} \times {}^{G}v_{g_z}\right)}{\left\| {}^{G}v_{OMC_y} \times {}^{G}v_{g_z} \right\|} \quad (1) $$

$$ {}^{G}v_{g_y} = {}^{G}v_{g_z} \times {}^{G}v_{g_x} \quad (2) $$

where $\times$ denotes the cross product. The orthonormal triad $\left\{ {}^{G}v_{g_x}, {}^{G}v_{g_y}, {}^{G}v_{g_z} \right\}$ was then converted to a quaternion ${}^{G}q_{g}$, reflecting the rotation mapping the original global OMC frame to the gravity-aligned global frame. All marker positions ${}^{G}p_{marker}$ originally expressed in ${}^{G}q_{OMC}$ were expressed in $g$ using ${}^{G}q_{g}$.

$$ {}^{g}p_{marker} = {}^{G}q_{g} \otimes {}^{G}p_{marker} \otimes \left({}^{G}q_{g}\right)^{*} \quad (3) $$

Where $(\quad)^{*}$ denotes the quaternion conjugate and $\otimes$ the quaternion multiplication.

Figure 3. Alignment of the OMC global Z-axis with the direction of gravity

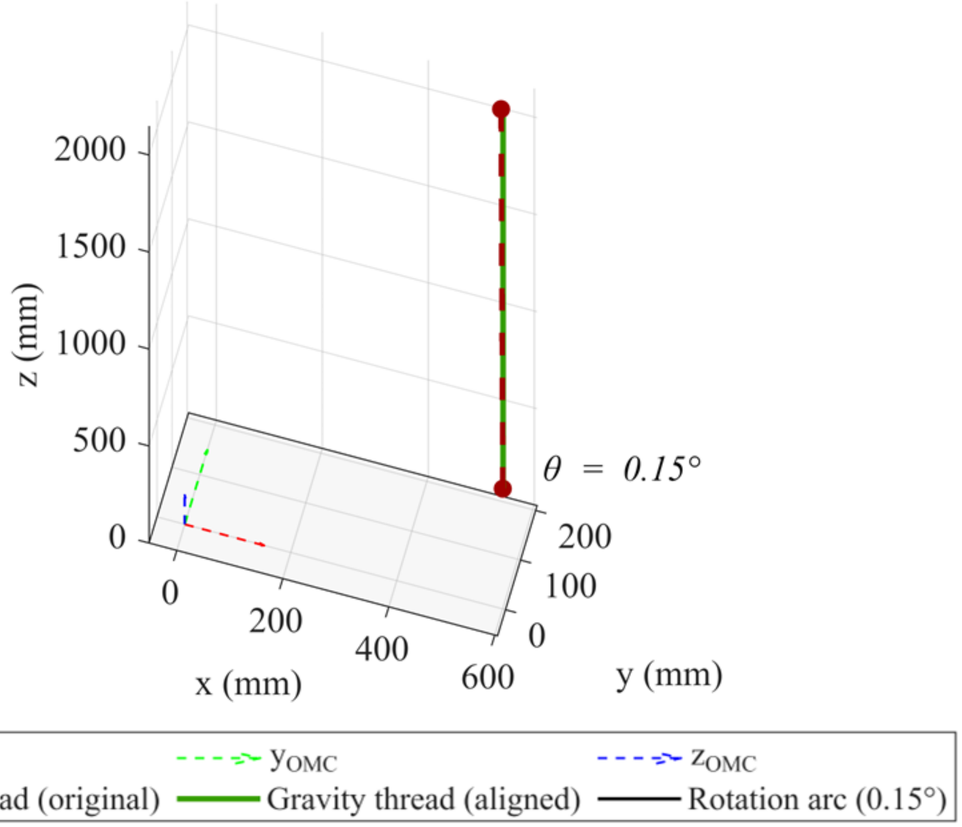


Legend: Vertical (gravity) vector estimated from the mean three-dimensional positions of reflective markers threaded through a line of a suspended pendulum within the OMC initial calibrated volume. The black arc denotes the rotation angle $\theta$ between the pre- and post-correction vertical unit vectors, computed as $\theta = cos^{-1}({}^{G}v_{OMC_z} \cdot {}^{G}v_{g_z})$ and used as a visual check. θ therefore represents the magnitude of the rotation removed by the gravity-alignment procedure and serves as a visual check that the OMC global Z-axis was correctly re-aligned with true vertical.

#### A.4. Misalignment between cluster and sensor frames

Alignment between the local IMU frames and cluster frames is essential for direct comparison of orientation estimates (Figure 4). Cluster-sensor misalignment may result from manual placement of the cluster relative to the sensor or slight discrepancies between the sensor frame and casing. A quaternion-based local frame alignment method (Lee & Jung, 2018) using gyroscope data and angular velocities derived from the marker cluster was used to estimate the rotational offset between the cluster frame and the attached IMU frame. Samples with low cluster angular velocity (< 0.2 rad/s on all three axes) were excluded prior to estimation, as such quasi-static periods provide poor directional information and can bias the optimisation. The offset quaternion was estimated by solving, in a least-squares sense, across the remaining samples k:

$$\omega_{IMU,k} = \left({}^{cluster}q_{IMU}\right)^{*} \otimes \omega_{OMC,k} \otimes {}^{cluster}q_{IMU} \tag{4}$$

where $\omega_{OMC,k}$ and $\omega_{IMU,k}$ are the cluster- and gyroscope-derived angular velocity vectors at sample k, expressed as pure quaternions, and ${}^{cluster}q_{IMU}$ is the sought rotation quaternion mapping the cluster frame onto the IMU frame. ${}^{cluster}q_{IMU}$ was obtained via constrained nonlinear least-squares (Levenberg–Marquardt algorithm):

$${}^{cluster}q_{IMU} = argmin_{q} \sum_{k=1}^{F} \left\| (q)^{*} \otimes \omega_{OMC,k} \otimes q - \omega_{IMU,k} \right\|^{2} \tag{5}$$

The estimated offset quaternion was then applied to correct the cluster orientation into the IMU-aligned frame:

$${}^{OMC}q_{cluster,aligned} = {}^{OMC}q_{cluster} \otimes {}^{cluster}q_{IMU} \tag{6}$$

Figure 4. Estimation of the rotational offset between the inertial measurement unit and rigid marker-cluster reference frames

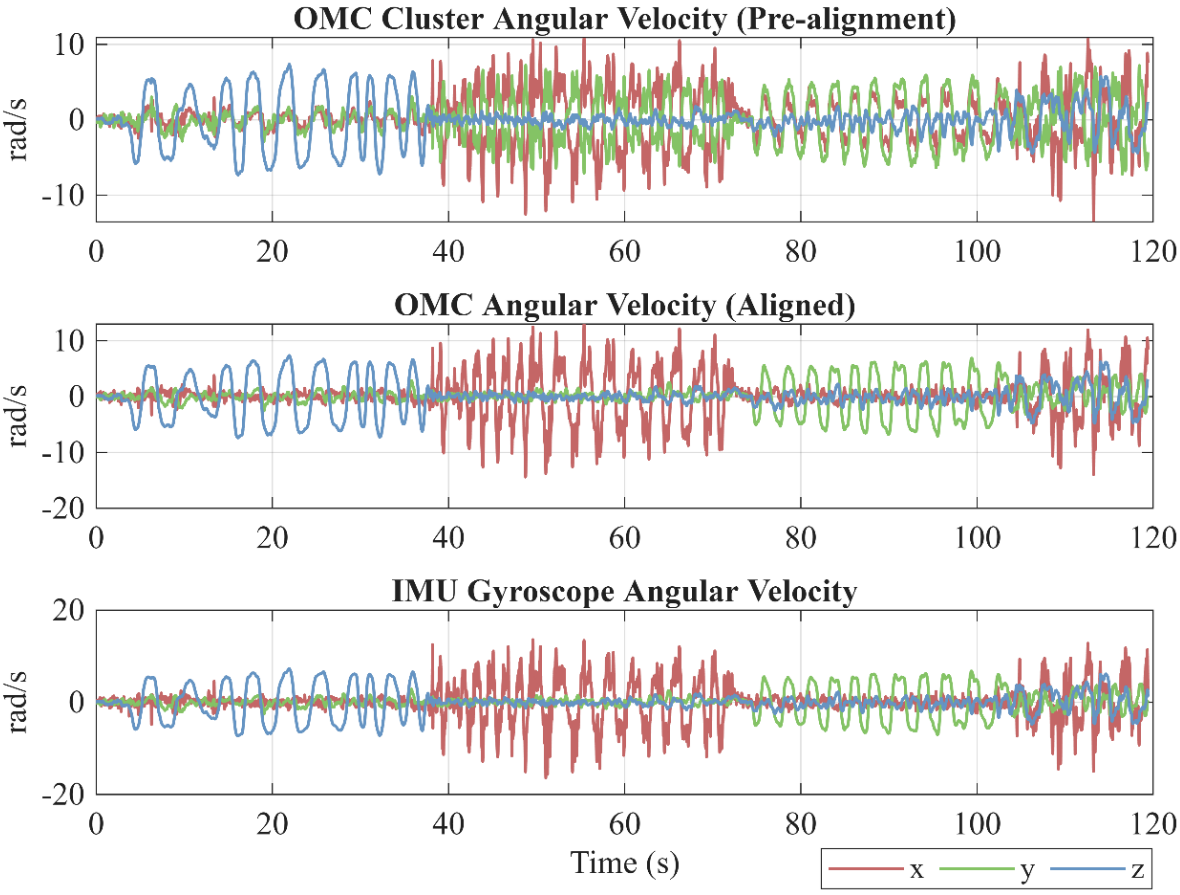


Legend: Representative angular velocity time series (x, y, z components; red, green, blue respectively) from a single participant and IMU/cluster unit. Top: cluster-derived angular velocity obtained from the numerical derivative of the OMC cluster orientation quaternion, expressed in the cluster frame. Middle: the same cluster-derived angular velocity after applying the estimated rotational offset quaternion, expressed in the IMU-aligned frame. Bottom: corresponding IMU gyroscope angular velocity.

#### A.5. Misalignment between global OMC x-axis and magnetic north

Ground truth cluster orientation was utilised to determine the magnetic north detected by the sensors' magnetometers during standing reference measures. To ensure accuracy, the magnetic field coordinates were expressed in the OMC frame, corrected for cluster orientation. Correction quaternions were then derived to rotate the OMC frame about the Z-axis, aligning it with the mean magnetometer values. This correction was applied to all marker measurements to adjust the heading (Figure 5).

Figure 5. Spherical representation of SLERP-interpolated rotations and magnetic heading vectors in the optical motion capture reference frame.

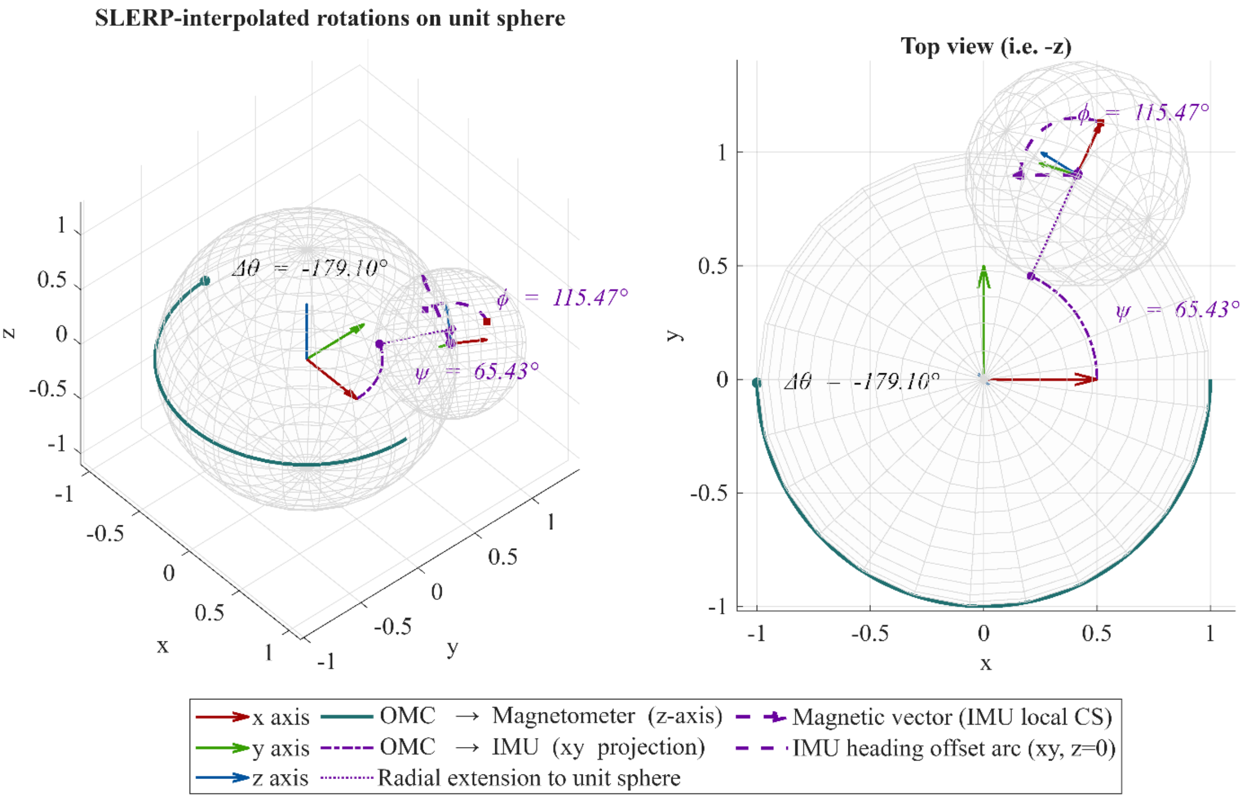


Legend: The central unit sphere represents the OMC reference frame. The solid teal arc illustrates the spherical linear interpolation trajectory of the rotation from the OMC frame toward the magnetometer heading direction (OMC → Magnetometer). The dash-dot purple arc depicts the projected rotation from the OMC frame toward the IMU x-axis local coordinate system (OMC → IMU), constrained to the horizontal plane ($z = 0$), with its angular extent annotated as $\psi$. The dotted purple line extending radially from the arc endpoint to the unit sphere surface indicates the direction of the IMU heading projected onto the global horizontal plane. The smaller sphere represents the IMU local reference frame, positioned at the endpoint of the OMC → IMU rotation and tilted to reflect the IMU orientation relative to the OMC global frame. The dashed purple arrow originating from the IMU sphere centre represents the magnetometer field vector expressed in the IMU local coordinate system. The dashed purple arc drawn around the IMU sphere centre subtends the angular offset $\varphi$ between the projected IMU x-axis and the projected magnetometer heading vector. The angular offset $\Delta\theta$ annotated at the teal arc endpoint denotes the total heading angle of the magnetometer heading vector relative to the OMC x-axis.

#### A.6. IMU downsampling

IMU orientation quaternions, obtained from the sensor-fusion filter at the native sampling rate of 1125 samples/s, were decimated by a factor of 9 to match the 125 samples/s sampling rate of the OMC system. Specifically, every ninth quaternion sample was retained rather than applying linear filtering, preserving the unit-norm constraint and avoiding the generation of potentially invalid intermediate orientations. Any residual discrepancy in sequence length was resolved by truncating the final quaternion sample.

### B. Sensor placement and Coordinate systems

#### B.1. IMU locations and alignment

Table 1. Manual alignment procedures with corresponding anatomical landmarks for each body segment.

| Location | Anatomical landmark | Sensor manual alignment |
|---|---|---|
| Thorax | Body of the sternum, below the sternal angle | Sensor x-axis aligned with the longitudinal axis of the thorax defined by the line connecting the processus xiphoideus (PX) to the incisura jugularis (IJ), pointing to PX. |
| Clavicle | Proximal shaft of the clavicle | Sensor x-axis aligned with the clavicular long axis defined by the line connecting the sternoclavicular joint (SC) to the acromioclavicular joint (AC), pointing toward AC. |
| Scapula | Angulus acromialis (AA) | Sensor x-axis aligned with the line connecting AA to AC, pointing toward AC. |
| Humerus | Distal humerus, dorsolateral surface at two-thirds of the arm length | Sensor x-axis aligned with the line connecting the glenohumeral joint centre (GHJC) and the midpoint between the medial (EM) and lateral (EL) epicondyles, pointing toward GHJC. |
| Forearm | Distal forearm, dorsal surface near the ulnar styloid | Sensor x-axis aligned with the line connecting the ulnar styloid (US) and the midpoint between EM and EL, pointing toward the midpoint. |
| Hand | Dorsal surface of the third metacarpal, distal region | Sensor x-axis aligned with the longitudinal axis of the third metacarpal, defined by the line connecting its head (MH3) and base (MB3), pointing toward the head. |

#### B.2. Estimation of the glenohumeral joint rotation centre

The glenohumeral joint rotation centre was estimated independently from the OMC and IMU data. For OMC, the joint centre was located using the Symmetrical Centre of Rotation Estimation (SCoRE) method (Ehrig et al., 2006), which finds the point whose coordinates, expressed in the local frames of the scapula and humerus segments, remain most consistent across the calibration trial without assuming either segment to be stationary.

For IMU, the joint centre was estimated from the scapula- and humerus-mounted sensors using (Figure 6) a two-segment formulation of the kinematic constraint proposed by Seel et al. (2012). Under rigid-body kinematics, the gravity-compensated specific force at any point on a segment can be related to the specific force measured by that segment's accelerometer through the segment's angular velocity $\omega$ and angular acceleration $\dot{\omega}$. Denoting by $r$ the constant, unknown position vector from an IMU to the joint centre, the specific force implied at the joint centre from segment $i$'s measurements is

$$a_{JC,i} = a_i - [\omega_i \times (\omega_i \times r_i) + \dot{\omega}_i \times r_i] = a_i - K_i\, r_i \qquad (7)$$

where the subscript $i \in \{1,2\}$ denotes the scapula and humerus segments, respectively; $a_i$ is the gravity-compensated specific force measured by segment $i$'s accelerometer; $r_i$ is the unknown, constant offset vector from that segment's IMU to the joint centre; and $K_i$ is the skew-symmetric matrix built from $\omega_i$ and $\dot{\omega}_i$ that maps $r_i$ onto the corresponding centripetal and tangential acceleration terms.

At the true joint centre, the specific force transported from each segment should coincide in magnitude. The offset vectors $r_1$ (scapula-to-joint centre) and $r_2$ (humerus-to-joint centre) are therefore obtained by minimising, in a nonlinear least-squares sense (Levenberg–Marquardt), the residual $\|a1 - K_1 r_1\| - \|a_2 - K_2 r_2\|$ across all calibration-movement samples.

Figure 6. Verification of the glenohumeral joint rotation centre estimated from IMU data using the symmetrical specific force centre method.

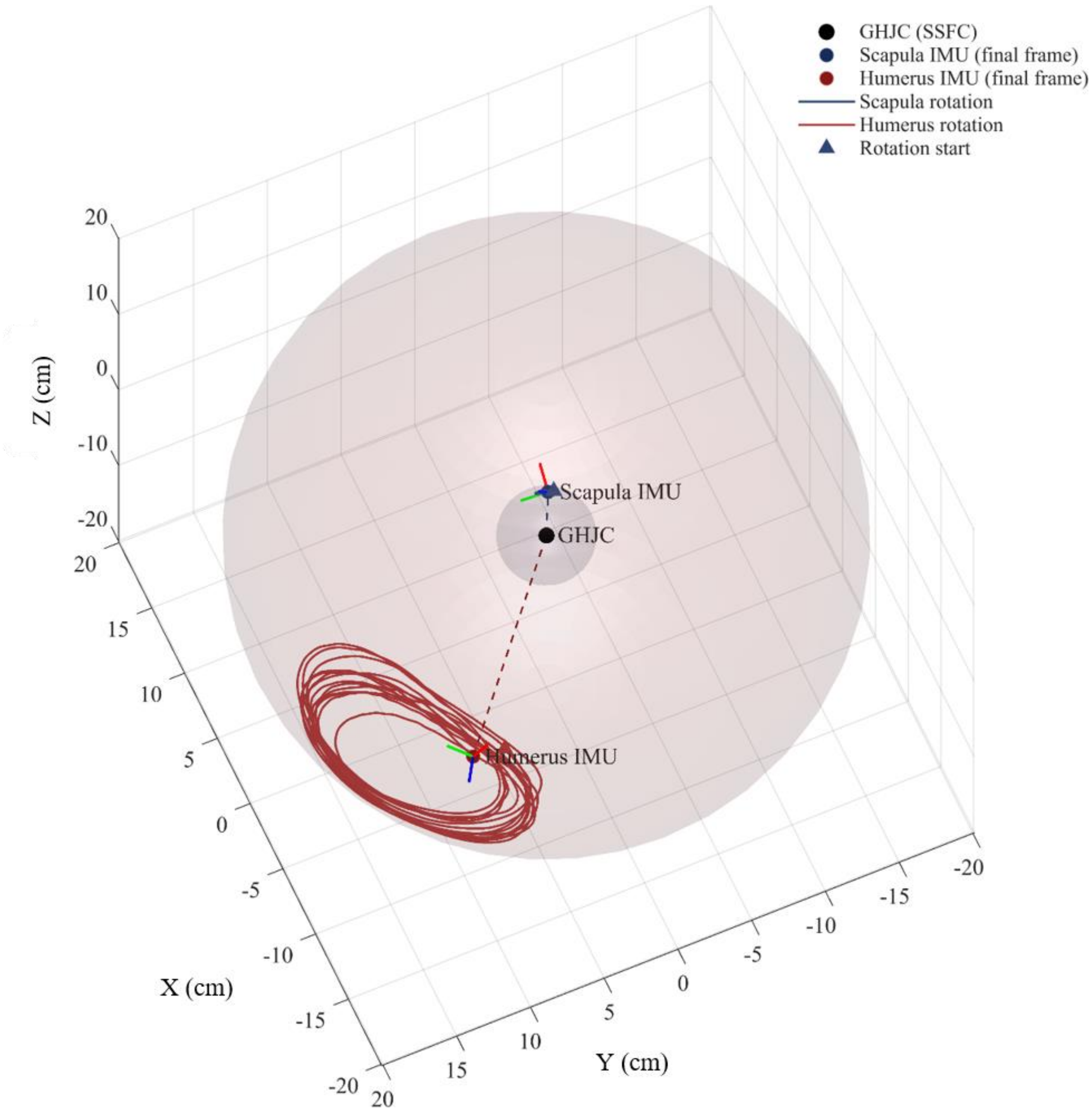


Legend: The black point marks the estimated glenohumeral joint rotation centre (GHJC) using the symmetrical specific force centre (SSFC) method. Blue and red points show the scapula and humerus IMU positions, respectively, at the final frame of the trial, connected to the GHJC by dashed offset vectors; red, green, and blue arrows at each IMU position denote its local x-, y-, and z-axes. The translucent blue and red spheres represent the theoretical position at a constant distance from the GHJC implied by the estimated scapula and humerus offset magnitudes. Solid blue and red curves show the scapula and humerus offset vectors, respectively, rotated into the global frame across the full calibration, with triangular markers indicating the start of each sweep.

#### B.3. Upper limb segment axes definition

Table 2. Definitions of segment coordinate systems and axis constructions for each body segment and alignment method.

| Segment | Origin | Axes | Axes definition | | | |
|---|---|---|---|---|---|---|
| | | | OMC* | IMU Neutral Alignment | IMU Manual Alignment | IMU Landmark Alignment |
| Thorax | $IJ$ | $x_T$ | $\widehat{Mid(PX,T8)\_IJ} \times \widehat{Mid(PX,T8)\_C7}$ | $IMU_x$ | $IMU_y$ | $\widehat{AC_L\_AC_R}$ |
| | | $y_T$ | $z_T \times x_T$ | $IMU_y$ | $IMU_z$ | $z_T \times x_T$ |
| | | $z_T$ | $\widehat{PX\_IJ + T8\_C7}$ | $IMU_z$ | $IMU_x$ | $\widehat{PX\_IJ + T8\_C7}$ |
| Clavicle | $SC$ | $x_C$ | $\widehat{SC\_AC}$ | $IMU_x$ | $IMU_x$ | $\widehat{SC\_AC}$ |
| | | $y_C$ | $z_T \times x_C$ | $IMU_y$ | $z_T \times x_C$ | $z_T \times x_C$ |
| | | $z_C$ | $x_C \times y_C$ | $IMU_z$ | $x_C \times y_C$ | $x_C \times y_C$ |
| Scapula | $AA$ | $x_S$ | $\widehat{TS\_AA}$ | $IMU_x$ | $-IMU_y$ | $\widehat{TS\_AA}$ |
| | | $y_S$ | $\widehat{IA\_TS} \times x_S$ | $IMU_y$ | $IMU_x$ | $\widehat{IA\_TS} \times x_S$ |
| | | $z_S$ | $x_S \times y_S$ | $IMU_z$ | $IMU_z$ | $x_S \times y_S$ |
| Humerus | $GH$ | $x_U$ | $y_U \times z_U$ | $IMU_x$ | $IMU_z$ | $y_U \times z_U$ |
| | | $y_U$ | $z_U \times \widehat{EM\_EL}$ | $IMU_y$ | $-IMU_y$ | $z_U \times \widehat{EM\_EL}$ |
| | | $z_U$ | $\widehat{Mid(EL,EM)\_GH}$ | $IMU_z$ | $IMU_x$ | $\widehat{Mid(EL,EM)\_GH}$ |
| Forearm | $US$ | $x_F$ | $y_F \times z_F$ | $IMU_x$ | $-IMU_y$ | $y_F \times z_F$ |
| | | $y_F$ | $z_F \times \widehat{US\_RS}$ | $IMU_y$ | $-IMU_z$ | $z_F \times \widehat{US\_RS}$ |
| | | $z_F$ | $\widehat{US\_Mid(EM,EL)}$ | $IMU_z$ | $IMU_x$ | $\widehat{US\_EM + US\_EL}$ |
| Hand | $M$ | $x_H$ | $y_H \times z_H$ | $IMU_x$ | $-IMU_y$ | $y_H \times z_H$ |
| | | $y_H$ | $z_H \times \widehat{US\_RS}$ | $IMU_y$ | $-IMU_z$ | $z_H \times \widehat{US\_RS}$ |
| | | $z_H$ | $\widehat{MH3\_MB3}$ | $IMU_z$ | $IMU_x$ | $\widehat{MH3\_MB3}$ |

Legend: $\hat{A}$ represents the unit vector $A$, $p_1\_p_2$ the vector from $p_1$ to $p_2$, $\times$ the cross product between two vectors, $Mid(p1,p2)$ the midpoint between $p_1$ and $p_2$ and $T, C, S, U, F, H$ are abbreviations for thorax, clavicle, scapula, humerus, forearm and hand, respectively. For IMU-based methods, axes labelled $IMU_x$, $IMU_y$, and $IMU_z$ correspond to the sensor-fixed coordinate system axes obtained either through landmark registration or neutral/manual alignment. *The International Society of Biomechanics (ISB)-recommended axes were permuted such that the original x-axis became the y-axis, the y-axis became the z-axis, and the z-axis became the x-axis.

#### B.4. Joint rotation sequences

Table 3. Joint definitions and Euler/Cardan rotation sequences used for joint angle decomposition according to ISB recommendations and modified conventions.

| **Joint** | **Parent** | **Child** | **ISB** | **Modified** |
|---|---|---|---|---|
| Sternoclavicular | Thorax | Clavicle | $YX'Z''$ | $ZY'X''$ |
| Scapulothoracic | Thorax | Scapula | $YX'Z''$ | $ZY'X''$ |
| Humerothoracic | Thorax | Humerus | $YX'Y''$ | $ZY'Z''$ |
| Acromioclavicular | Clavicle | Scapula | $YX'Z''$ | $ZY'X''$ |
| Glenohumeral | Scapula | Humerus | $YX'Y''$ | $YX'Z''$ |
| Elbow | Humerus | Forearm | $ZX'Y''$ | $XY'Z''$ |
| Wrist | Forearm | Hand | $YZ'X''$ | $XY'Z''$ |

Legend: Parent and Child denote the proximal and distal segments, respectively. The symbols $'$ and $''$ denote successive rotations about the axes of the rotating coordinate system after the first and second rotations, respectively.

### B.5. Scapula orientation estimation via the scapula-locator calibration method

A scapula locator was used to estimate three-dimensional scapular kinematics (Figure 7). With the participant positioned in a static reference posture, a three-pointed calliper-type instrument was manually aligned by the examiner onto the three ISB-recommended bony landmarks (acromial angle, trigonum spinae and inferior angle). The spatial arrangement of these landmarks is used to build the scapula's anatomical coordinate system, expressed as a unit quaternion in the global frame (${}^{G}q_{Sca}$). Simultaneously, the orientation of an IMU mounted over the acromion is sampled in the same global frame (${}^{G}q_{Acr}$). Because both quantities share a common frame and timestamp, the fixed rotational offset between ${}^{G}q_{Acr}$ and ${}^{G}q_{Sca}$ can be isolated:

$$ {}^{Acr}q_{Sca} = ({}^{G}q_{Acr})^{*} \otimes {}^{G}q_{Sca} \quad (8) $$

This offset is stored as a subject-specific calibration constant, representing the assumed-rigid relationship between the IMU and the scapula at the calibration pose. During the task, the locator is removed and only the acromion IMU remains. Scapula orientation is reconstructed at every sample by multiplying the live IMU orientation by the stored calibration offset:

$$ {}^{G}q_{Sca} = {}^{G}q_{Acr} \otimes {}^{Acr}q_{Sca} \quad (9) $$

Figure 7. Calibration and reconstruction of scapula orientation.

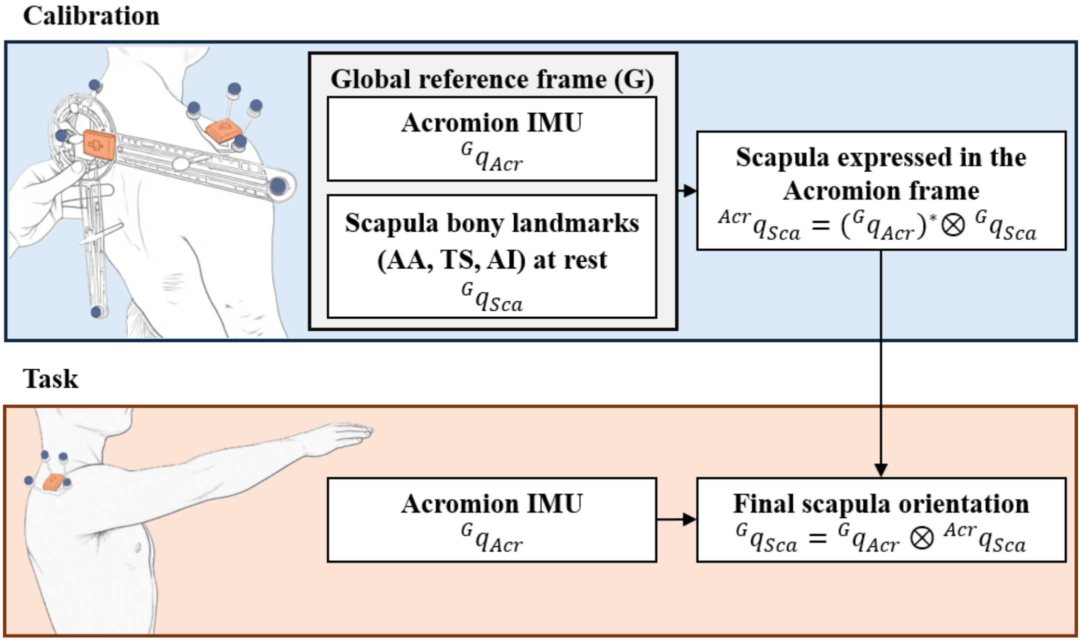


Legend: G, global reference frame; Acr, acromion; Sca, scapula; AA, TS, AI, Angulus Acromialis, Trigonum Spinae, Angulus Inferior; $q$, unit quaternion describing segment orientation; $\otimes$, quaternion product; *, quaternion conjugate.

### C. Analysis results

#### C.1. Exponential fit of gain versus angular velocity

Figure 8. Exponential mixed-effects modelling of sensor fusion gain as a function of segment angular velocity for each sensor type.

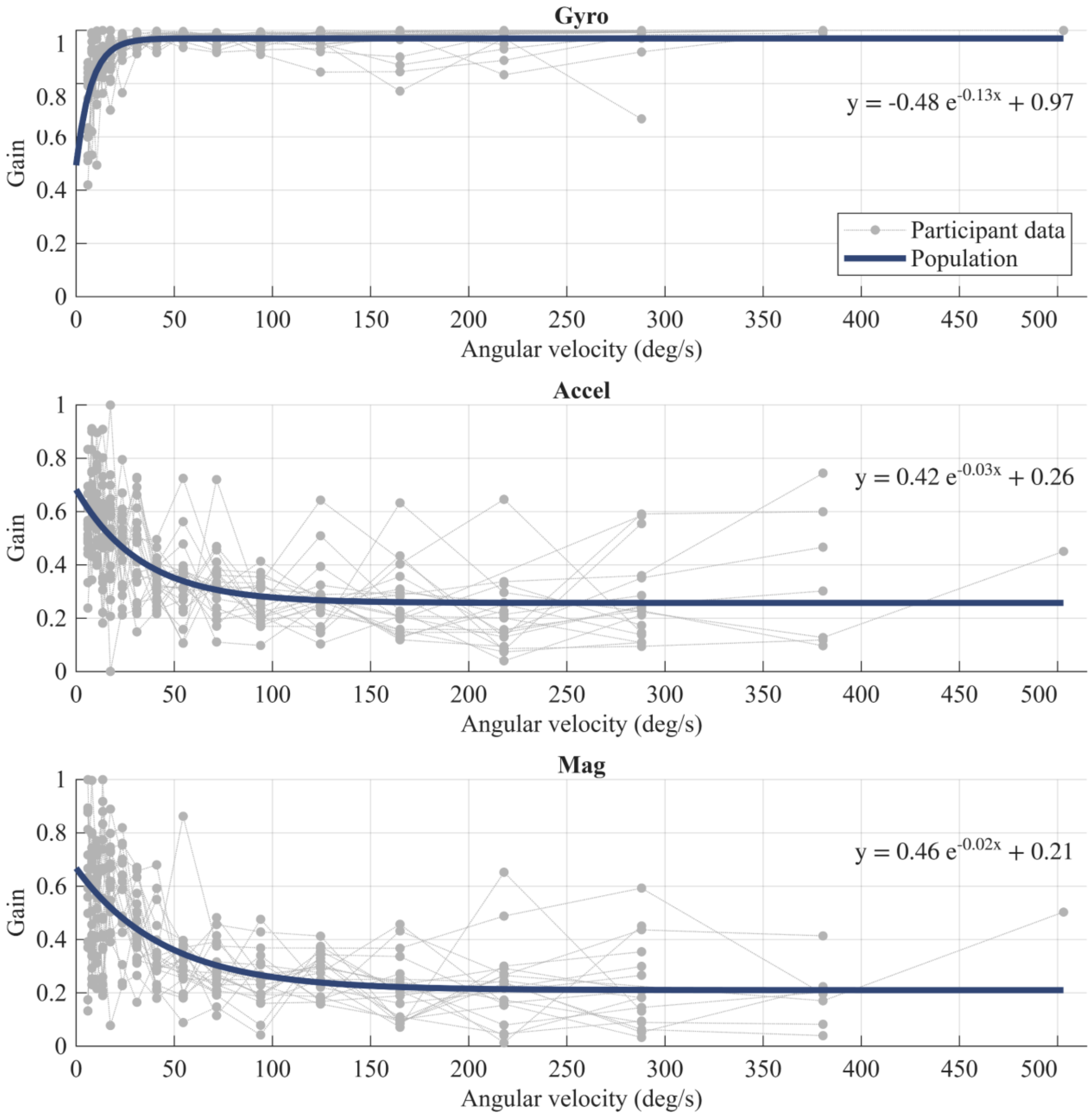


Legend: Grey dotted lines represent individual participant gain profiles across angular velocity bins, and the blue line corresponds to the population-level exponential fit.

#### C.2. Bland-Altman plots – Landmark method

For each joint–task combination, paired differences (OMC−IMU) and their corresponding mean angles were pooled across all participants into a single long-format dataset. A linear mixed-effects model was then fitted to this pooled dataset, with the OMC–IMU difference as the dependent variable, mean angle as a fixed-effect predictor, and a random intercept for participant ($LMM = DiffAngle \sim MeanAngle + (1 \mid Participant)$). The fixed-effect coefficient on mean angle characterises proportional bias — a significant non-zero slope indicates that the magnitude of the OMC–IMU discrepancy varies systematically with joint angle — while the model intercept reflects the fixed bias component at a mean angle of zero (Figure 9).

Figure 9. Bland-Altman comparison of joint angles between the landmark-based alignment and the OMC reference system.

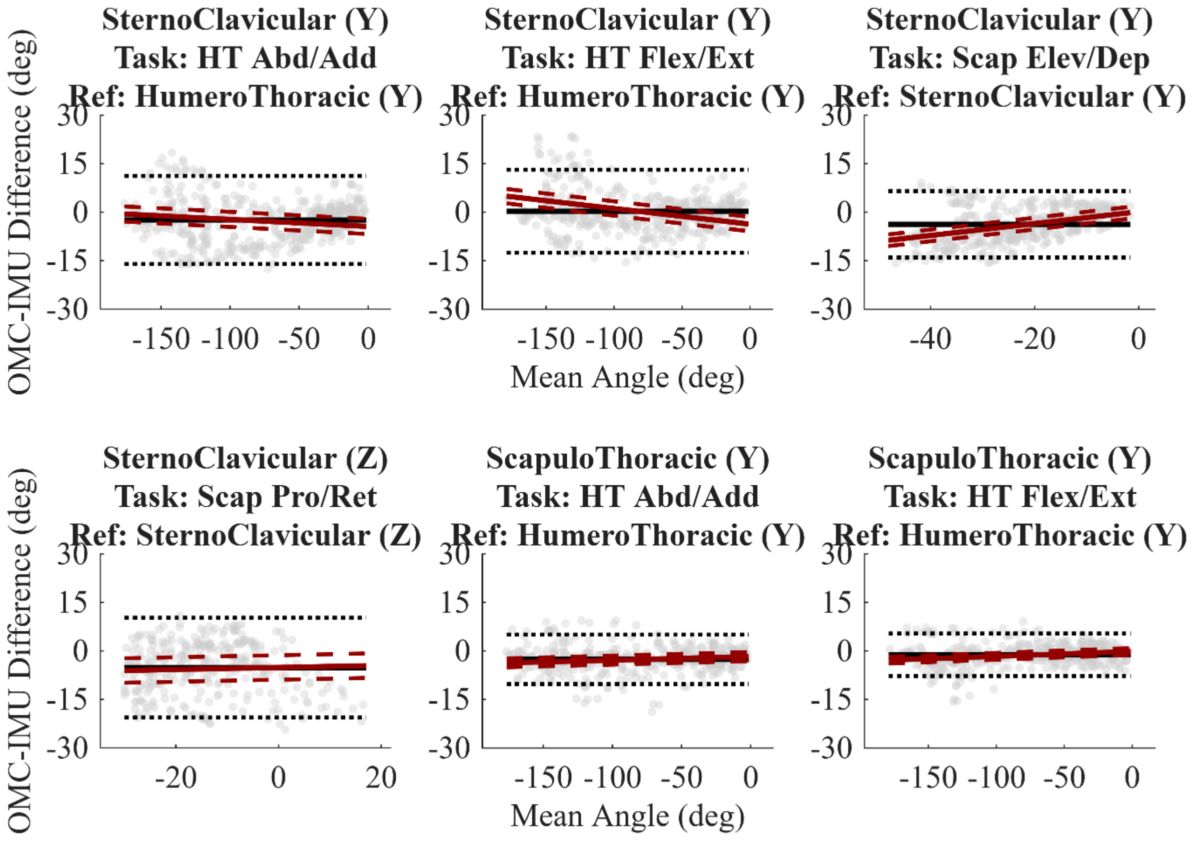

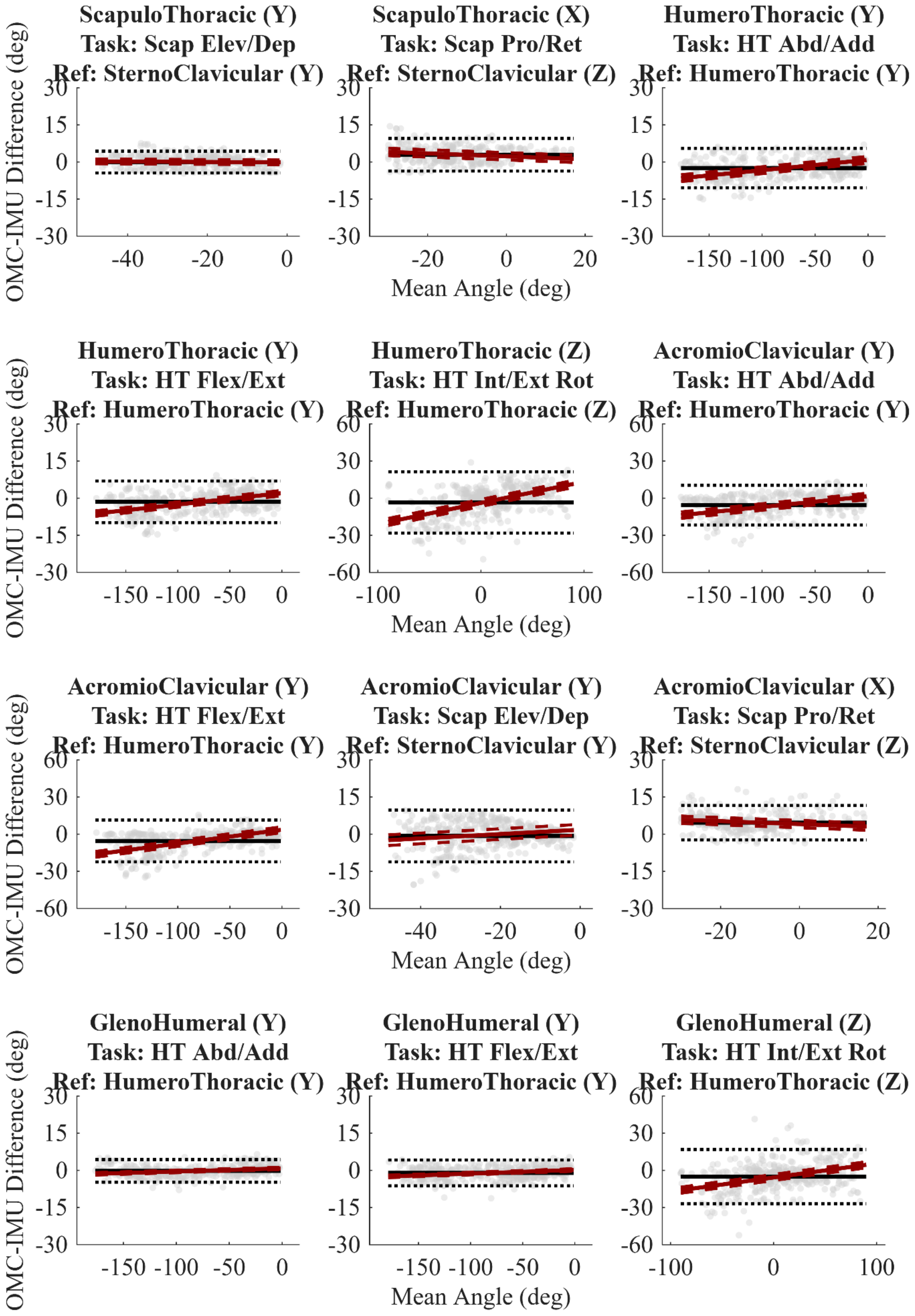
ScapuloThoracic (Y)
Task: Scap Elev/Dep
Ref: SternoClavicular (Y)
ScapuloThoracic (X)
Task: Scap Pro/Ret
Ref: SternoClavicular (Z)
HumeroThoracic (Y)
Task: HT Abd/Add
Ref: HumeroThoracic (Y)
HumeroThoracic (Y)
Task: HT Flex/Ext
Ref: HumeroThoracic (Y)
HumeroThoracic (Z)
Task: HT Int/Ext Rot
Ref: HumeroThoracic (Z)
AcromioClavicular (Y)
Task: HT Abd/Add
Ref: HumeroThoracic (Y)
AcromioClavicular (Y)
Task: HT Flex/Ext
Ref: HumeroThoracic (Y)
AcromioClavicular (Y)
Task: Scap Elev/Dep
Ref: SternoClavicular (Y)
AcromioClavicular (X)
Task: Scap Pro/Ret
Ref: SternoClavicular (Z)
GlenoHumeral (Y)
Task: HT Abd/Add
Ref: HumeroThoracic (Y)
GlenoHumeral (Y)
Task: HT Flex/Ext
Ref: HumeroThoracic (Y)
GlenoHumeral (Z)
Task: HT Int/Ext Rot
Ref: HumeroThoracic (Z)
OMC-IMU Difference (deg)
Mean Angle (deg)

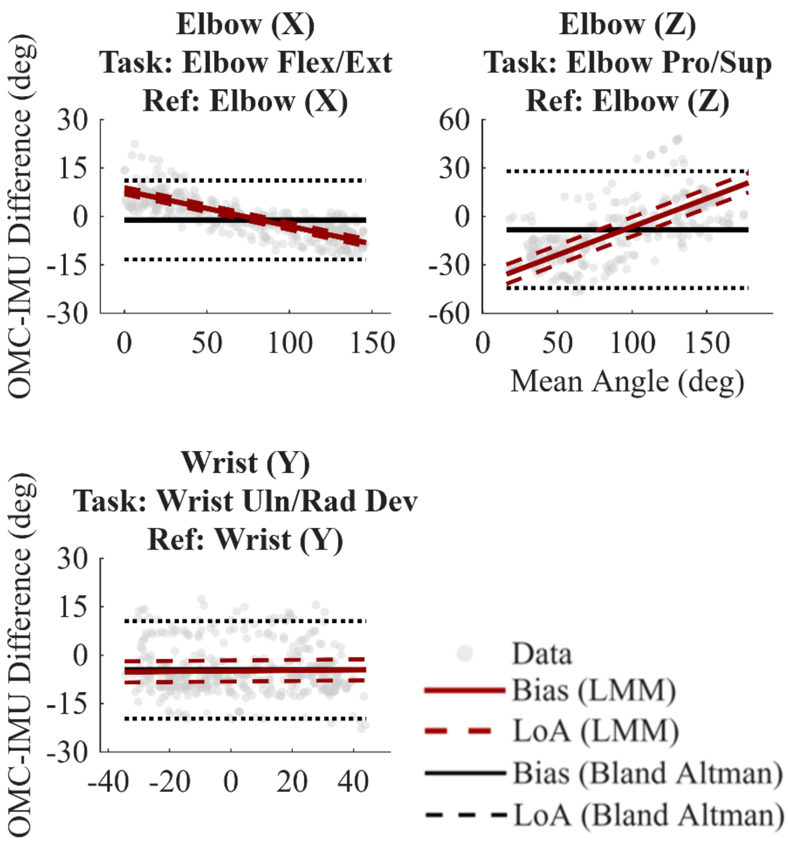


Legend: Grey scatter points represent individual frame-level observations downsampled for visualisation purposes, SC: sternoclavicular, AC: acromioclavicular, ST: scapulothoracic, HT: humerothoracic, GH: glenohumeral, EL: elbow, Pro/Sup: pronation/supination, Flex/Ext: flexion/extension, Scap Elev/Dep: scapular elevation depression, Abd/Add: abduction/adduction, Int/Ext Rot: internal/external rotation.

#### C.3. Bland-Altman parameters – Landmark method

Table 4. Bland-Altman and linear mixed-effects model results for IMU-derived versus OMC joint angle.

| Joint | Task | Bland-Altman Bias (LoA lower, LoA upper) | LMM Intercept (95% CI), p | LMM Coefficient °/° (95% CI), p |
|---|---|---|---|---|
| Sternoclavicular | HT Abd/Add | -2.4 (-16.0, 11.2) | -4.4 (-6.7, -2.1), p=0.0002 | -0.022 (-0.023, -0.021), p<0.0001 |
| Sternoclavicular | HT Flex/Ext | 0.3 (-12.5, 13.1) | -3.7 (-5.9, -1.5), p=0.001 | -0.048 (-0.049, -0.047), p<0.0001 |
| Sternoclavicular | Scap Elev/Dep | -3.7 (-14.0, 6.5) | 0.2 (-1.6, 2.0), p=0.86 | 0.184 (0.180, 0.187), p<0.0001 |
| Sternoclavicular | Scap Pro/Ret | -5.2 (-20.6, 10.3) | -5.1 (-8.9, -1.3), p=0.008 | 0.031 (0.026, 0.037), p<0.0001 |
| Scapulothoracic | HT Abd/Add | -2.6 (-10.3, 5.1) | -1.7 (-3.1, -0.3), p=0.014 | 0.011 (0.011, 0.012), p<0.0001 |
| Scapulothoracic | HT Flex/Ext | -1.2 (-7.8, 5.4) | -0.2 (-1.2, 0.9), p=0.765 | 0.015 (0.014, 0.015), p<0.0001 |
| Scapulothoracic | Scap Elev/Dep | -0.0 (-4.5, 4.4) | -0.1 (-1.1, 0.8), p=0.771 | -0.008 (-0.009, -0.007), p<0.0001 |
| Scapulothoracic | Scap Pro/Ret | 3.0 (-3.6, 9.6) | 2.3 (0.8, 3.8), p=0.002 | -0.059 (-0.062, -0.056), p<0.0001 |
| Humerothoracic | HT Abd/Add | -2.4 (-10.4, 5.5) | 0.9 (-0.5, 2.3), p=0.208 | 0.042 (0.042, 0.042), p<0.0001 |
| Humerothoracic | HT Flex/Ext | -1.5 (-9.9, 7.0) | 2.1 (1.0, 3.1), p=0.0002 | 0.046 (0.046, 0.047), p<0.0001 |
| Humerothoracic | HT Int/Ext Rota | -3.4 (-28.2, 21.4) | -3.8 (-6.6, -1.0), p=0.008 | 0.171 (0.169, 0.173), p<0.0001 |

| Joint | Task | Bland-Altman Bias (LoA lower, LoA upper) | LMM Intercept (95% CI), p | LMM Coefficient °/° (95% CI), p |
|---|---|---|---|---|
| Acromioclavicular | HT Abd/Add | -5.6 (-21.7, 10.5) | 1.5 (-1.1, 4.1), p=0.255 | 0.086 (0.085, 0.087), p<0.0001 |
| Acromioclavicular | HT Flex/Ext | -5.4 (-22.3, 11.4) | 3.6 (1.3, 5.9), p=0.002 | 0.111 (0.110, 0.112), p<0.0001 |
| Acromioclavicular | Scap Elev/Dep | -0.7 (-11.2, 9.7) | 1.8 (-0.3, 4.0), p=0.097 | 0.088 (0.084, 0.091), p<0.0001 |
| Acromioclavicular | Scap Pro/Ret | 4.6 (-2.3, 11.6) | 4.1 (2.6, 5.6), p<0.0001 | -0.059 (-0.063, -0.055), p<0.0001 |
| Glenohumeral | HT Abd/Add | -0.2 (-4.8, 4.4) | 0.8 (0.3, 1.3), p=0.001 | 0.013 (0.012, 0.013), p<0.0001 |
| Glenohumeral | HT Flex/Ext | -1.0 (-6.2, 4.2) | 0.2 (-0.3, 0.7), p=0.516 | 0.015 (0.014, 0.016), p<0.0001 |
| Glenohumeral | HT Int/Ext Rota | -5.0 (-26.9, 16.9) | -5.7 (-8.1, -3.3), p<0.0001 | 0.114 (0.112, 0.116), p<0.0001 |
| Elbow | Elbow Flex/Ext | -1.1 (-13.3, 11.1) | 8.0 (6.8, 9.1), p<0.0001 | -0.110 (-0.111, -0.109), p<0.0001 |
| Elbow | Elbow Pro/Sup | -8.2 (-44.3, 28.0) | -41.3 (-47.3, -35.3), p<0.0001 | 0.349 (0.347, 0.351), p<0.0001 |
| Wrist | Wrist Flex/Ext | -4.4 (-15.2, 6.4) | -4.7 (-6.5, -2.9), p<0.0001 | -0.036 (-0.036, -0.035), p<0.0001 |
| Wrist | Wrist Uln/Rad Dev | -4.5 (-19.6, 10.6) | -4.9 (-8.2, -1.6), p=0.003 | 0.008 (0.007, 0.009), p<0.0001 |

Legend: LoA, limits of agreement; LMM, linear mixed model; CI, confidence interval; HT, humerothoracic; Abd/Add, abduction/adduction; Flex/Ext, flexion/extension; Int/Ext Rota, internal/external rotation; Scap, scapular; Elev/Dep, elevation/depression; Pro/Ret, protraction/retraction; Uln/Rad Dev, ulnar/radial deviation; 95% CI = 95% confidence interval.

#### C.4. Joint*Movement interaction – Landmark method

Figure 10. Heatmap representation of the linear mixed model-derived EMMs across each joint*movement combination for the landmark alignment method.

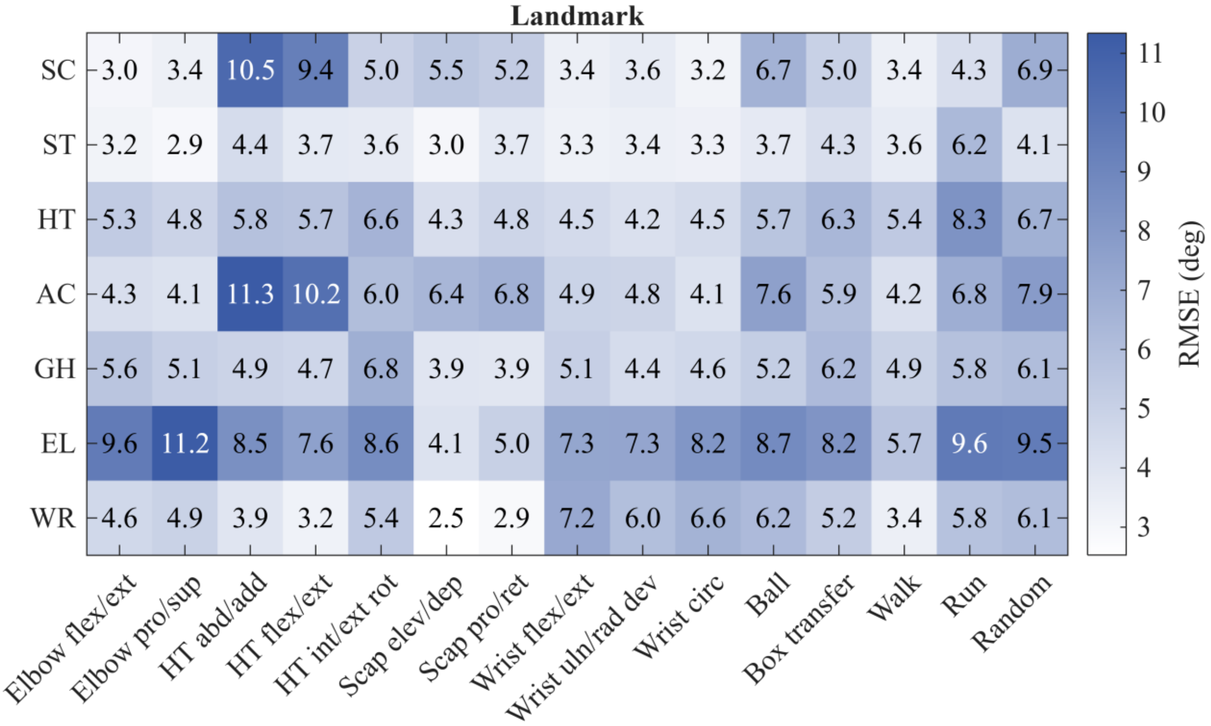


Legend: Legend: Flex/ext: flexion/extension, Pro/sup: protraction/supination, Abd/add: abduction/adduction, Int/ext rot: internal/external rotation, Scap: scapular, Elev/dep: elevation/depression, Pro/ret: protraction/retraction, Uln/rad dev: ulnar/radial deviation, , Circ: circumduction, SC: sternoclavicular, ST: scapulothoracic, AC: acromioclavicular, GH: Glenohumeral, HT: humerothoracic, EL: elbow, WR: wrist.

Table 5. Estimated marginal means of joint angle RMSE (°) for landmark alignment across all joint–movement combinations.

| Joint | Movement | EMM | SE | df | low.CL | upp.CL |
|---|---|---|---|---|---|---|
| Acromioclavicular | Elbow flex./ext. | 4.26 | 0.33 | 19 | 3.56 | 4.96 |
| Acromioclavicular | Elbow pro./sup. | 4.06 | 0.34 | 19 | 3.35 | 4.76 |
| Acromioclavicular | HT abd./add. | 11.34 | 0.33 | 19 | 10.64 | 12.03 |
| Acromioclavicular | HT flex./ext. | 10.17 | 0.33 | 19 | 9.47 | 10.86 |
| Acromioclavicular | HT int./ext. rot. | 6.01 | 0.33 | 19 | 5.31 | 6.7 |
| Acromioclavicular | HT rot. | 4.2 | 0.33 | 19 | 3.5 | 4.89 |

| Joint | Movement | EMM | SE | df | low.CL | upp.CL |
|---|---|---|---|---|---|---|
| Acromioclavicular | Shoulder girdle ele./dep. | 6.44 | 0.33 | 19 | 5.75 | 7.14 |
| Acromioclavicular | Shoulder girdle pro./ret. | 6.61 | 0.37 | 19 | 5.84 | 7.38 |
| Acromioclavicular | Wrist circ. | 4.08 | 0.33 | 19 | 3.38 | 4.77 |
| Acromioclavicular | Wrist uln./rad. dev. | 4.82 | 0.33 | 19 | 4.13 | 5.52 |
| Acromioclavicular | Wrist flex./ext. | 4.89 | 0.33 | 19 | 4.19 | 5.58 |
| Acromioclavicular | Ball | 7.59 | 0.33 | 19 | 6.89 | 8.29 |
| Acromioclavicular | Random | 7.86 | 0.51 | 19 | 6.79 | 8.93 |
| Acromioclavicular | Run | 6.84 | 0.34 | 19 | 6.13 | 7.55 |
| Acromioclavicular | Box transfer | 5.81 | 0.34 | 19 | 5.09 | 6.53 |
| Acromioclavicular | Walk | 4.18 | 0.33 | 19 | 3.48 | 4.87 |
| Elbow | Elbow flex./ext. | 9.57 | 0.38 | 19 | 8.78 | 10.36 |
| Elbow | Elbow pro./sup. | 11.23 | 0.38 | 19 | 10.42 | 12.03 |
| Elbow | HT abd./add. | 8.47 | 0.38 | 19 | 7.67 | 9.26 |
| Elbow | HT flex./ext. | 7.57 | 0.38 | 19 | 6.77 | 8.36 |
| Elbow | HT int./ext. rot. | 8.63 | 0.38 | 19 | 7.84 | 9.43 |
| Elbow | HT rot. | 5.22 | 0.38 | 19 | 4.43 | 6.02 |
| Elbow | Shoulder girdle ele./dep. | 4.08 | 0.38 | 19 | 3.29 | 4.88 |
| Elbow | Shoulder girdle pro./ret. | 4.87 | 0.42 | 19 | 3.98 | 5.76 |
| Elbow | Wrist circ. | 8.17 | 0.38 | 19 | 7.37 | 8.96 |
| Elbow | Wrist uln./rad. dev. | 7.34 | 0.38 | 19 | 6.54 | 8.13 |
| Elbow | Wrist flex./ext. | 7.29 | 0.38 | 19 | 6.5 | 8.09 |
| Elbow | Ball | 8.73 | 0.38 | 19 | 7.93 | 9.53 |
| Elbow | Random | 9.54 | 0.6 | 19 | 8.28 | 10.8 |
| Elbow | Run | 9.61 | 0.39 | 19 | 8.8 | 10.42 |
| Elbow | Box transfer | 8.11 | 0.4 | 19 | 7.29 | 8.94 |
| Elbow | Walk | 5.69 | 0.38 | 19 | 4.89 | 6.48 |
| Glenohumeral | Elbow flex./ext. | 5.61 | 0.28 | 19 | 5.02 | 6.2 |
| Glenohumeral | Elbow pro./sup. | 5.07 | 0.28 | 19 | 4.47 | 5.67 |
| Glenohumeral | HT abd./add. | 4.86 | 0.28 | 19 | 4.27 | 5.45 |
| Glenohumeral | HT flex./ext. | 4.71 | 0.28 | 19 | 4.12 | 5.3 |
| Glenohumeral | HT int./ext. rot. | 6.83 | 0.28 | 19 | 6.24 | 7.42 |
| Glenohumeral | HT rot. | 4.15 | 0.28 | 19 | 3.56 | 4.74 |
| Glenohumeral | Shoulder girdle ele./dep. | 3.94 | 0.28 | 19 | 3.35 | 4.53 |
| Glenohumeral | Shoulder girdle pro./ret. | 3.7 | 0.31 | 19 | 3.06 | 4.35 |
| Glenohumeral | Wrist circ. | 4.62 | 0.28 | 19 | 4.03 | 5.21 |
| Glenohumeral | Wrist uln./rad. dev. | 4.41 | 0.28 | 19 | 3.82 | 5 |
| Glenohumeral | Wrist flex./ext. | 5.1 | 0.28 | 19 | 4.51 | 5.69 |

| **Joint** | **Movement** | **EMM** | **SE** | **df** | **low.CL** | **upp.CL** |
|---|---|---|---|---|---|---|
| Glenohumeral | Ball | 5.14 | 0.28 | 19 | 4.55 | 5.74 |
| Glenohumeral | Random | 6.06 | 0.41 | 19 | 5.2 | 6.92 |
| Glenohumeral | Run | 5.84 | 0.29 | 19 | 5.24 | 6.44 |
| Glenohumeral | Box transfer | 6.08 | 0.29 | 19 | 5.47 | 6.69 |
| Glenohumeral | Walk | 4.93 | 0.28 | 19 | 4.34 | 5.52 |
| Humerothoracic | Elbow flex./ext. | 5.29 | 0.3 | 19 | 4.67 | 5.91 |
| Humerothoracic | Elbow pro./sup. | 4.82 | 0.3 | 19 | 4.19 | 5.44 |
| Humerothoracic | HT abd./add. | 5.8 | 0.3 | 19 | 5.18 | 6.42 |
| Humerothoracic | HT flex./ext. | 5.7 | 0.3 | 19 | 5.09 | 6.32 |
| Humerothoracic | HT int./ext. rot. | 6.58 | 0.3 | 19 | 5.96 | 7.2 |
| Humerothoracic | HT rot. | 3.88 | 0.3 | 19 | 3.26 | 4.5 |
| Humerothoracic | Shoulder girdle ele./dep. | 4.27 | 0.3 | 19 | 3.65 | 4.89 |
| Humerothoracic | Shoulder girdle pro./ret. | 4.64 | 0.32 | 19 | 3.96 | 5.32 |
| Humerothoracic | Wrist circ. | 4.45 | 0.3 | 19 | 3.83 | 5.07 |
| Humerothoracic | Wrist uln./rad. dev. | 4.15 | 0.3 | 19 | 3.53 | 4.76 |
| Humerothoracic | Wrist flex./ext. | 4.51 | 0.3 | 19 | 3.89 | 5.13 |
| Humerothoracic | Ball | 5.64 | 0.3 | 19 | 5.02 | 6.26 |
| Humerothoracic | Random | 6.71 | 0.44 | 19 | 5.79 | 7.63 |
| Humerothoracic | Run | 8.29 | 0.3 | 19 | 7.66 | 8.91 |
| Humerothoracic | Box transfer | 6.24 | 0.31 | 19 | 5.6 | 6.88 |
| Humerothoracic | Walk | 5.42 | 0.3 | 19 | 4.8 | 6.04 |
| Scapulothoracic | Elbow flex./ext. | 3.16 | 0.28 | 19 | 2.58 | 3.75 |
| Scapulothoracic | Elbow pro./sup. | 2.9 | 0.28 | 19 | 2.31 | 3.49 |
| Scapulothoracic | HT abd./add. | 4.4 | 0.28 | 19 | 3.82 | 4.98 |
| Scapulothoracic | HT flex./ext. | 3.72 | 0.28 | 19 | 3.14 | 4.3 |
| Scapulothoracic | HT int./ext. rot. | 3.65 | 0.28 | 19 | 3.06 | 4.23 |
| Scapulothoracic | HT rot. | 3.03 | 0.28 | 19 | 2.45 | 3.62 |
| Scapulothoracic | Shoulder girdle ele./dep. | 2.99 | 0.28 | 19 | 2.41 | 3.57 |
| Scapulothoracic | Shoulder girdle pro./ret. | 3.57 | 0.3 | 19 | 2.94 | 4.21 |
| Scapulothoracic | Wrist circ. | 3.29 | 0.28 | 19 | 2.7 | 3.87 |
| Scapulothoracic | Wrist uln./rad. dev. | 3.36 | 0.28 | 19 | 2.77 | 3.94 |
| Scapulothoracic | Wrist flex./ext. | 3.27 | 0.28 | 19 | 2.69 | 3.85 |
| Scapulothoracic | Ball | 3.68 | 0.28 | 19 | 3.1 | 4.27 |
| Scapulothoracic | Random | 4.1 | 0.4 | 19 | 3.26 | 4.95 |
| Scapulothoracic | Run | 6.25 | 0.28 | 19 | 5.65 | 6.83 |
| Scapulothoracic | Box transfer | 4.17 | 0.29 | 19 | 3.56 | 4.77 |
| Scapulothoracic | Walk | 3.61 | 0.28 | 19 | 3.02 | 4.19 |

| Joint | Movement | EMM | SE | df | low.CL | upp.CL |
|---|---|---|---|---|---|---|
| Sternoclavicular | Elbow flex./ext. | 3.03 | 0.34 | 19 | 2.32 | 3.75 |
| Sternoclavicular | Elbow pro./sup. | 3.33 | 0.35 | 19 | 2.6 | 4.05 |
| Sternoclavicular | HT abd./add. | 10.53 | 0.34 | 19 | 9.81 | 11.24 |
| Sternoclavicular | HT flex./ext. | 9.37 | 0.34 | 19 | 8.65 | 10.08 |
| Sternoclavicular | HT int./ext. rot. | 4.96 | 0.34 | 19 | 4.25 | 5.68 |
| Sternoclavicular | HT rot. | 2.71 | 0.34 | 19 | 2 | 3.43 |
| Sternoclavicular | Shoulder girdle ele./dep. | 5.5 | 0.34 | 19 | 4.78 | 6.21 |
| Sternoclavicular | Shoulder girdle pro./ret. | 5.09 | 0.38 | 19 | 4.29 | 5.88 |
| Sternoclavicular | Wrist circ. | 3.17 | 0.34 | 19 | 2.46 | 3.89 |
| Sternoclavicular | Wrist uln./rad. dev. | 3.57 | 0.34 | 19 | 2.85 | 4.29 |
| Sternoclavicular | Wrist flex./ext. | 3.39 | 0.34 | 19 | 2.67 | 4.1 |
| Sternoclavicular | Ball | 6.64 | 0.34 | 19 | 5.92 | 7.36 |
| Sternoclavicular | Random | 6.92 | 0.53 | 19 | 5.81 | 8.03 |
| Sternoclavicular | Run | 4.25 | 0.35 | 19 | 3.52 | 4.98 |
| Sternoclavicular | Box transfer | 4.87 | 0.36 | 19 | 4.12 | 5.61 |
| Sternoclavicular | Walk | 3.37 | 0.34 | 19 | 2.66 | 4.09 |
| Wrist | Elbow flex./ext. | 4.6 | 0.29 | 19 | 3.99 | 5.21 |
| Wrist | Elbow pro./sup. | 4.88 | 0.29 | 19 | 4.27 | 5.5 |
| Wrist | HT abd./add. | 3.94 | 0.29 | 19 | 3.33 | 4.55 |
| Wrist | HT flex./ext. | 3.25 | 0.29 | 19 | 2.64 | 3.86 |
| Wrist | HT int./ext. rot. | 5.41 | 0.29 | 19 | 4.8 | 6.01 |
| Wrist | HT rot. | 3.17 | 0.29 | 19 | 2.56 | 3.78 |
| Wrist | Shoulder girdle ele./dep. | 2.53 | 0.29 | 19 | 1.92 | 3.14 |
| Wrist | Shoulder girdle pro./ret. | 2.71 | 0.32 | 19 | 2.05 | 3.38 |
| Wrist | Wrist circ. | 6.62 | 0.29 | 19 | 6.01 | 7.23 |
| Wrist | Wrist uln./rad. dev. | 5.97 | 0.29 | 19 | 5.36 | 6.58 |
| Wrist | Wrist flex./ext. | 7.17 | 0.29 | 19 | 6.56 | 7.77 |
| Wrist | Ball | 6.21 | 0.29 | 19 | 5.59 | 6.82 |
| Wrist | Random | 6.06 | 0.43 | 19 | 5.16 | 6.96 |
| Wrist | Run | 5.76 | 0.29 | 19 | 5.14 | 6.38 |
| Wrist | Box transfer | 5.13 | 0.3 | 19 | 4.5 | 5.76 |
| Wrist | Walk | 3.36 | 0.29 | 19 | 2.75 | 3.96 |

Legend: EMM: estimated marginal mean, SE: standard error, df: degrees of freedom, low.CL: lower confidence limit, upp.CL: upper confidence limit, HT: humerothoracic; ele./dep.: elevation/depression; pro./ret.: protraction/retraction; abd./add.: abduction/adduction;

flex./ext.: flexion/extension; int./ext. rot.: internal/external rotation; uln./rad. dev.: ulnar/radial deviation; circ.: circumduction, low: lower, up: upper, CL: confidence limits.

#### C.5. Joint*Movement interaction – Manual method

Figure 11. Heatmap representation of the linear mixed model-derived EMMs across each joint*movement combination for the manual alignment method.

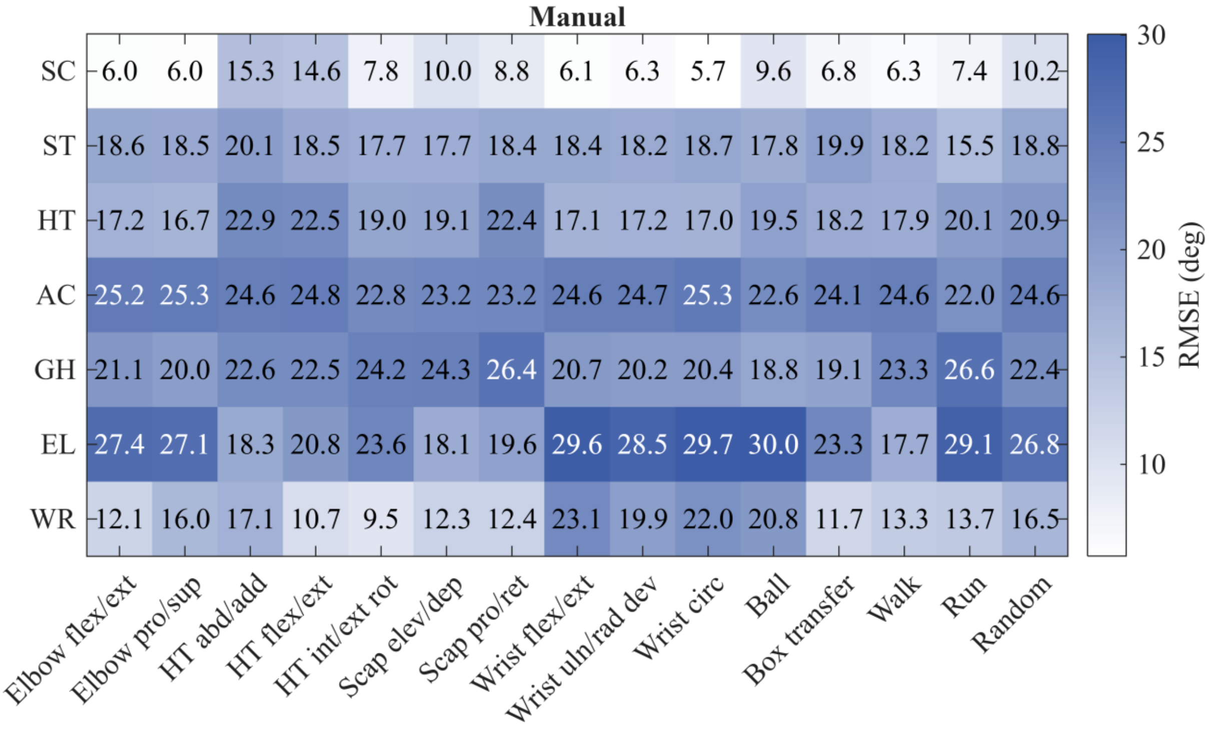


Legend: Legend: Flex/ext: flexion/extension, Pro/sup: protraction/supination, Abd/add: abduction/adduction, Int/ext rot: internal/external rotation, Scap: scapular, Elev/dep: elevation/depression, Pro/ret: protraction/retraction, Uln/rad dev: ulnar/radial deviation, , Circ: circumduction, SC: sternoclavicular, ST: scapulothoracic, AC: acromioclavicular, GH: Glenohumeral, HT: humerothoracic, EL: elbow, WR: wrist.

Table 6. Estimated marginal means of joint angle RMSE (°) for manual alignment across all joint–movement combinations.

| **Joint** | **Movement** | **EMM** | **SE** | **df** | **low.CL** | **upp.CL** |
|---|---|---|---|---|---|---|
| Acromioclavicular | Elbow flex./ext. | 25.18 | 0.65 | 19 | 23.83 | 26.54 |
| Acromioclavicular | Elbow pro./sup. | 25.24 | 0.65 | 19 | 23.88 | 26.61 |
| Acromioclavicular | HT abd./add. | 24.65 | 0.65 | 19 | 23.30 | 26.01 |
| Acromioclavicular | HT flex./ext. | 24.76 | 0.65 | 19 | 23.40 | 26.11 |
| Acromioclavicular | HT int./ext. rot. | 22.77 | 0.65 | 19 | 21.41 | 24.12 |
| Acromioclavicular | HT rot. | 23.85 | 0.65 | 19 | 22.50 | 25.21 |
| Acromioclavicular | Shoulder girdle ele./dep. | 23.20 | 0.65 | 19 | 21.84 | 24.55 |

| Joint | Movement | EMM | SE | df | low.CL | upp.CL |
|---|---|---|---|---|---|---|
| Acromioclavicular | Shoulder girdle pro./ret. | 23.18 | 0.71 | 19 | 21.70 | 24.66 |
| Acromioclavicular | Wrist circ. | 25.31 | 0.65 | 19 | 23.95 | 26.66 |
| Acromioclavicular | Wrist uln./rad. dev. | 24.66 | 0.65 | 19 | 23.31 | 26.02 |
| Acromioclavicular | Wrist flex./ext. | 24.55 | 0.65 | 19 | 23.20 | 25.90 |
| Acromioclavicular | Ball | 22.62 | 0.65 | 19 | 21.25 | 23.98 |
| Acromioclavicular | Random | 24.56 | 0.95 | 19 | 22.57 | 26.54 |
| Acromioclavicular | Run | 22.06 | 0.66 | 19 | 20.68 | 23.44 |
| Acromioclavicular | Box transfer | 24.29 | 0.67 | 19 | 22.89 | 25.68 |
| Acromioclavicular | Walk | 24.61 | 0.65 | 19 | 23.26 | 25.96 |
| Elbow | Elbow flex./ext. | 27.45 | 0.92 | 19 | 25.51 | 29.38 |
| Elbow | Elbow pro./sup. | 27.02 | 0.94 | 19 | 25.06 | 28.98 |
| Elbow | HT abd./add. | 18.31 | 0.92 | 19 | 16.37 | 20.24 |
| Elbow | HT flex./ext. | 20.78 | 0.92 | 19 | 18.85 | 22.71 |
| Elbow | HT int./ext. rot. | 23.57 | 0.92 | 19 | 21.64 | 25.50 |
| Elbow | HT rot. | 18.26 | 0.92 | 19 | 16.33 | 20.19 |
| Elbow | Shoulder girdle ele./dep. | 18.13 | 0.92 | 19 | 16.19 | 20.06 |
| Elbow | Shoulder girdle pro./ret. | 19.49 | 1.04 | 19 | 17.32 | 21.67 |
| Elbow | Wrist circ. | 29.67 | 0.92 | 19 | 27.74 | 31.61 |
| Elbow | Wrist uln./rad. dev. | 28.53 | 0.92 | 19 | 26.60 | 30.46 |
| Elbow | Wrist flex./ext. | 29.58 | 0.92 | 19 | 27.65 | 31.51 |
| Elbow | Ball | 30.03 | 0.93 | 19 | 28.08 | 31.98 |
| Elbow | Random | 26.78 | 1.49 | 19 | 23.68 | 29.89 |
| Elbow | Run | 29.09 | 0.94 | 19 | 27.12 | 31.06 |
| Elbow | Box transfer | 23.46 | 0.96 | 19 | 21.45 | 25.48 |
| Elbow | Walk | 17.70 | 0.92 | 19 | 15.76 | 19.63 |
| Glenohumeral | Elbow flex./ext. | 21.07 | 0.93 | 19 | 19.12 | 23.02 |
| Glenohumeral | Elbow pro./sup. | 19.97 | 0.94 | 19 | 18.00 | 21.94 |
| Glenohumeral | HT abd./add. | 22.60 | 0.93 | 19 | 20.65 | 24.55 |
| Glenohumeral | HT flex./ext. | 22.47 | 0.93 | 19 | 20.52 | 24.41 |
| Glenohumeral | HT int./ext. rot. | 24.24 | 0.93 | 19 | 22.30 | 26.19 |
| Glenohumeral | HT rot. | 21.41 | 0.93 | 19 | 19.47 | 23.36 |
| Glenohumeral | Shoulder girdle ele./dep. | 24.31 | 0.93 | 19 | 22.37 | 26.26 |
| Glenohumeral | Shoulder girdle pro./ret. | 26.34 | 1.05 | 19 | 24.15 | 28.53 |
| Glenohumeral | Wrist circ. | 20.39 | 0.93 | 19 | 18.44 | 22.33 |
| Glenohumeral | Wrist uln./rad. dev. | 20.22 | 0.93 | 19 | 18.27 | 22.17 |
| Glenohumeral | Wrist flex./ext. | 20.73 | 0.93 | 19 | 18.79 | 22.68 |
| Glenohumeral | Ball | 18.76 | 0.94 | 19 | 16.80 | 20.72 |

| Joint | Movement | EMM | SE | df | low.CL | upp.CL |
|---|---|---|---|---|---|---|
| Glenohumeral | Random | 22.35 | 1.50 | 19 | 19.22 | 25.48 |
| Glenohumeral | Run | 26.70 | 0.95 | 19 | 24.71 | 28.68 |
| Glenohumeral | Box transfer | 19.34 | 0.97 | 19 | 17.31 | 21.37 |
| Glenohumeral | Walk | 23.29 | 0.93 | 19 | 21.34 | 25.23 |
| Humerothoracic | Elbow flex./ext. | 17.18 | 0.64 | 19 | 15.85 | 18.52 |
| Humerothoracic | Elbow pro./sup. | 16.65 | 0.64 | 19 | 15.31 | 18.00 |
| Humerothoracic | HT abd./add. | 22.88 | 0.64 | 19 | 21.55 | 24.22 |
| Humerothoracic | HT flex./ext. | 22.47 | 0.64 | 19 | 21.14 | 23.81 |
| Humerothoracic | HT int./ext. rot. | 19.02 | 0.64 | 19 | 17.69 | 20.35 |
| Humerothoracic | HT rot. | 18.40 | 0.64 | 19 | 17.07 | 19.74 |
| Humerothoracic | Shoulder girdle ele./dep. | 19.07 | 0.64 | 19 | 17.74 | 20.40 |
| Humerothoracic | Shoulder girdle pro./ret. | 22.28 | 0.69 | 19 | 20.83 | 23.74 |
| Humerothoracic | Wrist circ. | 16.99 | 0.64 | 19 | 15.66 | 18.33 |
| Humerothoracic | Wrist uln./rad. dev. | 17.19 | 0.64 | 19 | 15.86 | 18.52 |
| Humerothoracic | Wrist flex./ext. | 17.10 | 0.64 | 19 | 15.77 | 18.43 |
| Humerothoracic | Ball | 19.49 | 0.64 | 19 | 18.15 | 20.83 |
| Humerothoracic | Random | 20.86 | 0.93 | 19 | 18.92 | 22.81 |
| Humerothoracic | Run | 20.09 | 0.65 | 19 | 18.74 | 21.44 |
| Humerothoracic | Box transfer | 18.42 | 0.66 | 19 | 17.04 | 19.79 |
| Humerothoracic | Walk | 17.93 | 0.64 | 19 | 16.60 | 19.26 |
| Scapulothoracic | Elbow flex./ext. | 18.58 | 0.73 | 19 | 17.06 | 20.11 |
| Scapulothoracic | Elbow pro./sup. | 18.47 | 0.74 | 19 | 16.93 | 20.01 |
| Scapulothoracic | HT abd./add. | 20.11 | 0.73 | 19 | 18.59 | 21.63 |
| Scapulothoracic | HT flex./ext. | 18.51 | 0.73 | 19 | 16.99 | 20.04 |
| Scapulothoracic | HT int./ext. rot. | 17.70 | 0.73 | 19 | 16.18 | 19.23 |
| Scapulothoracic | HT rot. | 18.71 | 0.73 | 19 | 17.19 | 20.24 |
| Scapulothoracic | Shoulder girdle ele./dep. | 17.73 | 0.73 | 19 | 16.21 | 19.25 |
| Scapulothoracic | Shoulder girdle pro./ret. | 18.29 | 0.80 | 19 | 16.61 | 19.98 |
| Scapulothoracic | Wrist circ. | 18.71 | 0.73 | 19 | 17.18 | 20.23 |
| Scapulothoracic | Wrist uln./rad. dev. | 18.22 | 0.73 | 19 | 16.70 | 19.74 |
| Scapulothoracic | Wrist flex./ext. | 18.36 | 0.73 | 19 | 16.84 | 19.88 |
| Scapulothoracic | Ball | 17.80 | 0.73 | 19 | 16.27 | 19.33 |
| Scapulothoracic | Random | 18.82 | 1.11 | 19 | 16.50 | 21.15 |
| Scapulothoracic | Run | 15.57 | 0.74 | 19 | 14.02 | 17.12 |
| Scapulothoracic | Box transfer | 20.06 | 0.75 | 19 | 18.49 | 21.64 |
| Scapulothoracic | Walk | 18.21 | 0.73 | 19 | 16.69 | 19.73 |
| Sternoclavicular | Elbow flex./ext. | 6.03 | 0.61 | 19 | 4.74 | 7.32 |

| Joint | Movement | EMM | SE | df | low.CL | upp.CL |
|---|---|---|---|---|---|---|
| Sternoclavicular | Elbow pro./sup. | 5.92 | 0.62 | 19 | 4.62 | 7.22 |
| Sternoclavicular | HT abd./add. | 15.31 | 0.61 | 19 | 14.02 | 16.59 |
| Sternoclavicular | HT flex./ext. | 14.58 | 0.61 | 19 | 13.29 | 15.87 |
| Sternoclavicular | HT int./ext. rot. | 7.80 | 0.61 | 19 | 6.52 | 9.09 |
| Sternoclavicular | HT rot. | 5.82 | 0.61 | 19 | 4.54 | 7.11 |
| Sternoclavicular | Shoulder girdle ele./dep. | 10.01 | 0.61 | 19 | 8.73 | 11.30 |
| Sternoclavicular | Shoulder girdle pro./ret. | 8.78 | 0.67 | 19 | 7.38 | 10.18 |
| Sternoclavicular | Wrist circ. | 5.73 | 0.61 | 19 | 4.45 | 7.02 |
| Sternoclavicular | Wrist uln./rad. dev. | 6.29 | 0.61 | 19 | 5.00 | 7.57 |
| Sternoclavicular | Wrist flex./ext. | 6.06 | 0.61 | 19 | 4.78 | 7.35 |
| Sternoclavicular | Ball | 9.56 | 0.62 | 19 | 8.27 | 10.86 |
| Sternoclavicular | Random | 10.25 | 0.88 | 19 | 8.40 | 12.10 |
| Sternoclavicular | Run | 7.43 | 0.62 | 19 | 6.12 | 8.73 |
| Sternoclavicular | Box transfer | 6.97 | 0.63 | 19 | 5.65 | 8.30 |
| Sternoclavicular | Walk | 6.32 | 0.61 | 19 | 5.03 | 7.60 |
| Wrist | Elbow flex./ext. | 12.08 | 0.68 | 19 | 10.65 | 13.51 |
| Wrist | Elbow pro./sup. | 16.00 | 0.69 | 19 | 14.56 | 17.44 |
| Wrist | HT abd./add. | 17.13 | 0.68 | 19 | 15.71 | 18.56 |
| Wrist | HT flex./ext. | 10.73 | 0.68 | 19 | 9.30 | 12.15 |
| Wrist | HT int./ext. rot. | 9.51 | 0.68 | 19 | 8.08 | 10.94 |
| Wrist | HT rot. | 14.62 | 0.68 | 19 | 13.20 | 16.05 |
| Wrist | Shoulder girdle ele./dep. | 12.34 | 0.68 | 19 | 10.91 | 13.77 |
| Wrist | Shoulder girdle pro./ret. | 12.30 | 0.75 | 19 | 10.73 | 13.87 |
| Wrist | Wrist circ. | 22.03 | 0.68 | 19 | 20.60 | 23.46 |
| Wrist | Wrist uln./rad. dev. | 19.90 | 0.68 | 19 | 18.47 | 21.33 |
| Wrist | Wrist flex./ext. | 23.07 | 0.68 | 19 | 21.65 | 24.50 |
| Wrist | Ball | 20.75 | 0.69 | 19 | 19.32 | 22.19 |
| Wrist | Random | 16.51 | 1.02 | 19 | 14.37 | 18.64 |
| Wrist | Run | 13.71 | 0.69 | 19 | 12.26 | 15.16 |
| Wrist | Box transfer | 11.89 | 0.71 | 19 | 10.41 | 13.36 |
| Wrist | Walk | 13.31 | 0.68 | 19 | 11.88 | 14.74 |

Legend: EMM: estimated marginal mean, SE: standard error, df: degrees of freedom, low.CL: lower confidence limit, upp.CL: upper confidence limit, HT: humerothoracic; ele./dep.: elevation/depression; pro./ret.: protraction/retraction; abd./add.: abduction/adduction; flex./ext.: flexion/extension; int./ext. rot.: internal/external rotation; uln./rad. dev.: ulnar/radial deviation; circ.: circumduction, low: lower, up: upper, CL: confidence limits.

#### C.6. Joint*Movement interaction – Neutral method

Figure 12. Heatmap representation of the linear mixed model-derived EMMs across each joint*movement combination for the neutral alignment method.

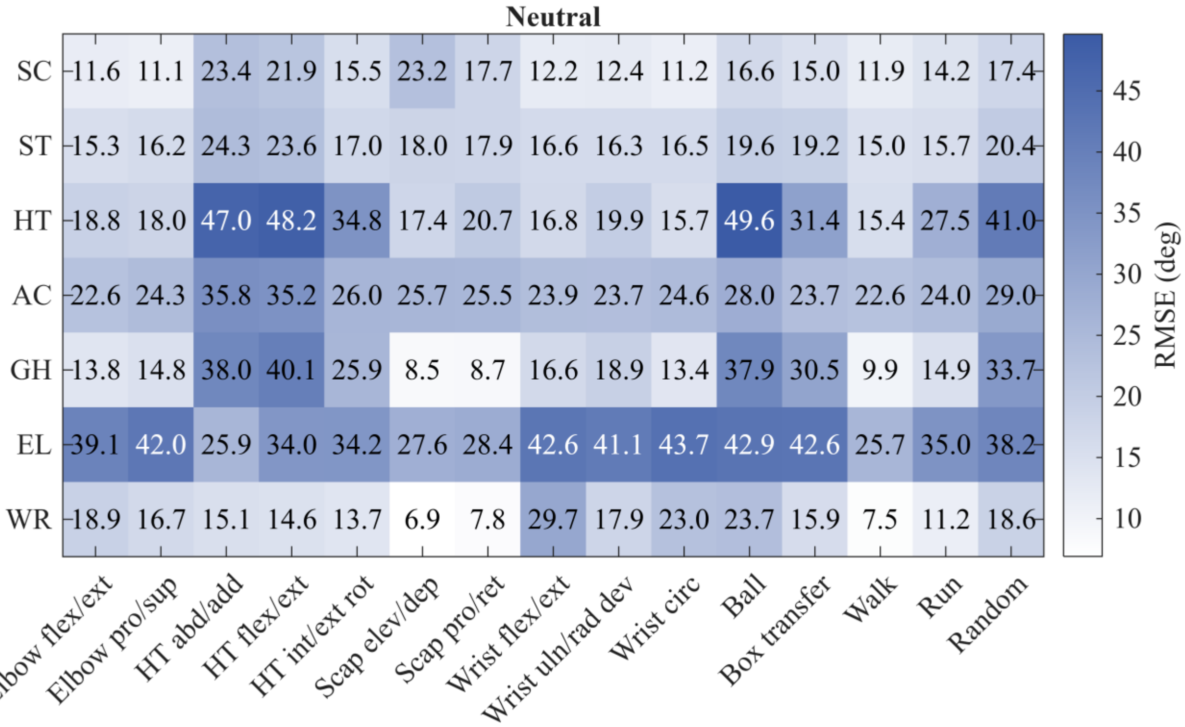


Legend: Legend: Flex/ext: flexion/extension, Pro/sup: protraction/supination, Abd/add: abduction/adduction, Int/ext rot: internal/external rotation, Scap: scapular, Elev/dep: elevation/depression, Pro/ret: protraction/retraction, Uln/rad dev: ulnar/radial deviation, , Circ: circumduction, SC: sternoclavicular, ST: scapulothoracic, AC: acromioclavicular, GH: Glenohumeral, HT: humerothoracic, EL: elbow, WR: wrist.

Table 7. Estimated marginal means of joint angle RMSE (°) for neutral alignment across all joint–movement combinations.

| Joint | Movement | EMM | SE | df | low.CL | upp.CL |
|---|---|---|---|---|---|---|
| Acromioclavicular | Elbow flex./ext. | 22.62 | 0.67 | 19 | 21.23 | 24.02 |
| Acromioclavicular | Elbow pro./sup. | 24.29 | 0.67 | 19 | 22.88 | 25.70 |
| Acromioclavicular | HT abd./add. | 35.79 | 0.67 | 19 | 34.39 | 37.18 |
| Acromioclavicular | HT flex./ext. | 35.21 | 0.67 | 19 | 33.82 | 36.61 |
| Acromioclavicular | HT int./ext. rot. | 26.01 | 0.67 | 19 | 24.61 | 27.40 |
| Acromioclavicular | HT rot. | 23.42 | 0.67 | 19 | 22.03 | 24.82 |
| Acromioclavicular | Shoulder girdle ele./dep. | 25.74 | 0.67 | 19 | 24.35 | 27.13 |
| Acromioclavicular | Shoulder girdle pro./ret. | 25.68 | 0.72 | 19 | 24.18 | 27.19 |

| Joint | Movement | EMM | SE | df | low.CL | upp.CL |
|---|---|---|---|---|---|---|
| Acromioclavicular | Wrist circ. | 24.60 | 0.67 | 19 | 23.21 | 26.00 |
| Acromioclavicular | Wrist uln./rad. dev. | 23.70 | 0.67 | 19 | 22.30 | 25.09 |
| Acromioclavicular | Wrist flex./ext. | 23.92 | 0.67 | 19 | 22.52 | 25.31 |
| Acromioclavicular | Ball | 28.00 | 0.67 | 19 | 26.60 | 29.40 |
| Acromioclavicular | Random | 28.99 | 0.94 | 19 | 27.03 | 30.95 |
| Acromioclavicular | Run | 23.92 | 0.67 | 19 | 22.51 | 25.33 |
| Acromioclavicular | Box transfer | 23.73 | 0.68 | 19 | 22.30 | 25.16 |
| Acromioclavicular | Walk | 22.65 | 0.67 | 19 | 21.25 | 24.04 |
| Elbow | Elbow flex./ext. | 39.06 | 1.64 | 19 | 35.63 | 42.49 |
| Elbow | Elbow pro./sup. | 42.01 | 1.66 | 19 | 38.53 | 45.49 |
| Elbow | HT abd./add. | 25.85 | 1.64 | 19 | 22.43 | 29.28 |
| Elbow | HT flex./ext. | 33.97 | 1.64 | 19 | 30.54 | 37.40 |
| Elbow | HT int./ext. rot. | 34.22 | 1.64 | 19 | 30.79 | 37.64 |
| Elbow | HT rot. | 25.37 | 1.64 | 19 | 21.94 | 28.79 |
| Elbow | Shoulder girdle ele./dep. | 27.60 | 1.64 | 19 | 24.18 | 31.03 |
| Elbow | Shoulder girdle pro./ret. | 28.61 | 1.87 | 19 | 24.69 | 32.52 |
| Elbow | Wrist circ. | 43.70 | 1.64 | 19 | 40.27 | 47.12 |
| Elbow | Wrist uln./rad. dev. | 41.13 | 1.64 | 19 | 37.70 | 44.55 |
| Elbow | Wrist flex./ext. | 42.57 | 1.64 | 19 | 39.14 | 45.99 |
| Elbow | Ball | 42.92 | 1.65 | 19 | 39.47 | 46.37 |
| Elbow | Random | 38.24 | 2.75 | 19 | 32.48 | 44.01 |
| Elbow | Run | 34.89 | 1.68 | 19 | 31.38 | 38.40 |
| Elbow | Box transfer | 42.70 | 1.72 | 19 | 39.10 | 46.30 |
| Elbow | Walk | 25.67 | 1.64 | 19 | 22.24 | 29.09 |
| Glenohumeral | Elbow flex./ext. | 13.75 | 1.40 | 19 | 10.82 | 16.69 |
| Glenohumeral | Elbow pro./sup. | 14.86 | 1.42 | 19 | 11.88 | 17.83 |
| Glenohumeral | HT abd./add. | 37.95 | 1.40 | 19 | 35.02 | 40.88 |
| Glenohumeral | HT flex./ext. | 40.11 | 1.40 | 19 | 37.18 | 43.05 |
| Glenohumeral | HT int./ext. rot. | 25.86 | 1.40 | 19 | 22.93 | 28.80 |
| Glenohumeral | HT rot. | 19.58 | 1.40 | 19 | 16.65 | 22.51 |
| Glenohumeral | Shoulder girdle ele./dep. | 8.49 | 1.40 | 19 | 5.56 | 11.43 |
| Glenohumeral | Shoulder girdle pro./ret. | 8.88 | 1.60 | 19 | 5.54 | 12.22 |
| Glenohumeral | Wrist circ. | 13.45 | 1.40 | 19 | 10.52 | 16.39 |
| Glenohumeral | Wrist uln./rad. dev. | 18.94 | 1.40 | 19 | 16.01 | 21.87 |
| Glenohumeral | Wrist flex./ext. | 16.64 | 1.40 | 19 | 13.70 | 19.57 |
| Glenohumeral | Ball | 37.90 | 1.41 | 19 | 34.94 | 40.85 |
| Glenohumeral | Random | 33.69 | 2.33 | 19 | 28.81 | 38.57 |

| Joint | Movement | EMM | SE | df | low.CL | upp.CL |
|---|---|---|---|---|---|---|
| Glenohumeral | Run | 14.84 | 1.43 | 19 | 11.84 | 17.84 |
| Glenohumeral | Box transfer | 30.55 | 1.47 | 19 | 27.48 | 33.63 |
| Glenohumeral | Walk | 9.86 | 1.40 | 19 | 6.93 | 12.80 |
| Humerothoracic | Elbow flex./ext. | 18.82 | 1.37 | 19 | 15.96 | 21.68 |
| Humerothoracic | Elbow pro./sup. | 17.99 | 1.39 | 19 | 15.09 | 20.90 |
| Humerothoracic | HT abd./add. | 47.03 | 1.37 | 19 | 44.17 | 49.89 |
| Humerothoracic | HT flex./ext. | 48.19 | 1.37 | 19 | 45.33 | 51.05 |
| Humerothoracic | HT int./ext. rot. | 34.80 | 1.37 | 19 | 31.94 | 37.66 |
| Humerothoracic | HT rot. | 25.54 | 1.37 | 19 | 22.68 | 28.40 |
| Humerothoracic | Shoulder girdle ele./dep. | 17.44 | 1.37 | 19 | 14.58 | 20.30 |
| Humerothoracic | Shoulder girdle pro./ret. | 20.93 | 1.55 | 19 | 17.67 | 24.18 |
| Humerothoracic | Wrist circ. | 15.68 | 1.37 | 19 | 12.82 | 18.54 |
| Humerothoracic | Wrist uln./rad. dev. | 19.87 | 1.37 | 19 | 17.01 | 22.73 |
| Humerothoracic | Wrist flex./ext. | 16.75 | 1.37 | 19 | 13.88 | 19.61 |
| Humerothoracic | Ball | 49.63 | 1.38 | 19 | 46.75 | 52.51 |
| Humerothoracic | Random | 40.99 | 2.27 | 19 | 36.24 | 45.74 |
| Humerothoracic | Run | 27.41 | 1.40 | 19 | 24.48 | 30.33 |
| Humerothoracic | Box transfer | 31.42 | 1.43 | 19 | 28.43 | 34.42 |
| Humerothoracic | Walk | 15.45 | 1.37 | 19 | 12.59 | 18.31 |
| Scapulothoracic | Elbow flex./ext. | 15.32 | 0.64 | 19 | 13.97 | 16.67 |
| Scapulothoracic | Elbow pro./sup. | 16.20 | 0.65 | 19 | 14.84 | 17.55 |
| Scapulothoracic | HT abd./add. | 24.27 | 0.64 | 19 | 22.93 | 25.62 |
| Scapulothoracic | HT flex./ext. | 23.55 | 0.64 | 19 | 22.20 | 24.90 |
| Scapulothoracic | HT int./ext. rot. | 16.97 | 0.64 | 19 | 15.63 | 18.32 |
| Scapulothoracic | HT rot. | 16.15 | 0.64 | 19 | 14.80 | 17.50 |
| Scapulothoracic | Shoulder girdle ele./dep. | 18.02 | 0.64 | 19 | 16.67 | 19.37 |
| Scapulothoracic | Shoulder girdle pro./ret. | 18.09 | 0.69 | 19 | 16.64 | 19.54 |
| Scapulothoracic | Wrist circ. | 16.49 | 0.64 | 19 | 15.14 | 17.84 |
| Scapulothoracic | Wrist uln./rad. dev. | 16.34 | 0.64 | 19 | 14.99 | 17.69 |
| Scapulothoracic | Wrist flex./ext. | 16.58 | 0.64 | 19 | 15.23 | 17.93 |
| Scapulothoracic | Ball | 19.61 | 0.65 | 19 | 18.26 | 20.96 |
| Scapulothoracic | Random | 20.35 | 0.89 | 19 | 18.50 | 22.21 |
| Scapulothoracic | Run | 15.67 | 0.65 | 19 | 14.31 | 17.03 |
| Scapulothoracic | Box transfer | 19.30 | 0.66 | 19 | 17.92 | 20.68 |
| Scapulothoracic | Walk | 15.03 | 0.64 | 19 | 13.68 | 16.38 |
| Sternoclavicular | Elbow flex./ext. | 11.64 | 0.79 | 19 | 10.00 | 13.28 |
| Sternoclavicular | Elbow pro./sup. | 11.06 | 0.79 | 19 | 9.40 | 12.73 |

| Joint | Movement | EMM | SE | df | low.CL | upp.CL |
|---|---|---|---|---|---|---|
| Sternoclavicular | HT abd./add. | 23.37 | 0.79 | 19 | 21.73 | 25.01 |
| Sternoclavicular | HT flex./ext. | 21.95 | 0.79 | 19 | 20.30 | 23.59 |
| Sternoclavicular | HT int./ext. rot. | 15.50 | 0.79 | 19 | 13.85 | 17.14 |
| Sternoclavicular | HT rot. | 11.81 | 0.79 | 19 | 10.17 | 13.46 |
| Sternoclavicular | Shoulder girdle ele./dep. | 23.19 | 0.79 | 19 | 21.55 | 24.83 |
| Sternoclavicular | Shoulder girdle pro./ret. | 17.91 | 0.86 | 19 | 16.10 | 19.72 |
| Sternoclavicular | Wrist circ. | 11.19 | 0.79 | 19 | 9.54 | 12.83 |
| Sternoclavicular | Wrist uln./rad. dev. | 12.39 | 0.79 | 19 | 10.74 | 14.03 |
| Sternoclavicular | Wrist flex./ext. | 12.17 | 0.79 | 19 | 10.53 | 13.82 |
| Sternoclavicular | Ball | 16.62 | 0.79 | 19 | 14.96 | 18.27 |
| Sternoclavicular | Random | 17.45 | 1.18 | 19 | 14.98 | 19.93 |
| Sternoclavicular | Run | 14.17 | 0.80 | 19 | 12.50 | 15.85 |
| Sternoclavicular | Box transfer | 15.04 | 0.81 | 19 | 13.34 | 16.74 |
| Sternoclavicular | Walk | 11.89 | 0.79 | 19 | 10.25 | 13.53 |
| Wrist | Elbow flex./ext. | 18.92 | 0.91 | 19 | 17.02 | 20.82 |
| Wrist | Elbow pro./sup. | 16.67 | 0.92 | 19 | 14.74 | 18.60 |
| Wrist | HT abd./add. | 15.12 | 0.91 | 19 | 13.22 | 17.02 |
| Wrist | HT flex./ext. | 14.64 | 0.91 | 19 | 12.73 | 16.54 |
| Wrist | HT int./ext. rot. | 13.70 | 0.91 | 19 | 11.79 | 15.60 |
| Wrist | HT rot. | 8.72 | 0.91 | 19 | 6.82 | 10.63 |
| Wrist | Shoulder girdle ele./dep. | 6.91 | 0.91 | 19 | 5.01 | 8.81 |
| Wrist | Shoulder girdle pro./ret. | 8.04 | 1.01 | 19 | 5.92 | 10.17 |
| Wrist | Wrist circ. | 23.02 | 0.91 | 19 | 21.12 | 24.92 |
| Wrist | Wrist uln./rad. dev. | 17.88 | 0.91 | 19 | 15.98 | 19.78 |
| Wrist | Wrist flex./ext. | 29.69 | 0.91 | 19 | 27.79 | 31.59 |
| Wrist | Ball | 23.69 | 0.91 | 19 | 21.77 | 25.60 |
| Wrist | Random | 18.57 | 1.42 | 19 | 15.59 | 21.55 |
| Wrist | Run | 11.14 | 0.93 | 19 | 9.20 | 13.08 |
| Wrist | Box transfer | 15.94 | 0.95 | 19 | 13.96 | 17.92 |
| Wrist | Walk | 7.51 | 0.91 | 19 | 5.61 | 9.41 |

Legend: EMM: estimated marginal mean, SE: standard error, df: degrees of freedom, low.CL: lower confidence limit, upp.CL: upper confidence limit, HT: humerothoracic; ele./dep.: elevation/depression; pro./ret.: protraction/retraction; abd./add.: abduction/adduction; flex./ext.: flexion/extension; int./ext. rot.: internal/external rotation; uln./rad. dev.: ulnar/radial deviation; circ.: circumduction, low: lower, up: upper, CL: confidence limits.

### D. References

Ehrig, R. M., Taylor, W. R., Duda, G. N., & Heller, M. O. (2006). A survey of formal methods for determining the centre of rotation of ball joints. *J Biomech*, *39*(15), 2798-2809. https://doi.org/10.1016/j.jbiomech.2005.10.002

Lee, J. K., & Jung, W. C. (2018). Quaternion-based local frame alignment between an inertial measurement unit and a motion capture system. *Sensors (Basel)*, *18*(11). https://doi.org/10.3390/s18114003

Seel, T., Schauer, T., & Raisch, J. (2012). *Joint axis and position estimation from inertial measurement data by exploiting kinematic constraints* 2012 IEEE International Conference on Control Applications,
